# Multiscale computer model of the spinal dorsal horn reveals changes in network processing associated with chronic pain

**DOI:** 10.1101/2021.06.09.447785

**Authors:** Laura Medlock, Kazutaka Sekiguchi, Sungho Hong, Salvador Dura-Bernal, William W Lytton, Steven A. Prescott

**Affiliations:** Neurosciences and Mental Health, The Hospital for Sick Children, Toronto, Ontario M5G 0A4, Canada; Institute of Biomedical Engineering, University of Toronto, Toronto, Ontario M5S 3G9, Canada; Drug Developmental Research Laboratory, Shionogi Pharmaceutical Research Center, Toyonaka, Osaka 561-0825, Japan; State University of New York Downstate Health Science University, Brooklyn, NY 11203, US; Computational Neuroscience Unit, Okinawa Institute of Science and Technology, Okinawa 904-0495, Japan; Nathan Kline Institute for Psychiatric Research, Orangeburg, NY 10962, US; Kings County Hospital, Brooklyn, NY 11207, US; Department of Physiology, University of Toronto, Toronto, Ontario M5S 1A8, Canada

**Author notes:** LM and KS contributed equally to this research. SAP and WWL contributed equally to this research. To whom correspondence should be addressed: Steven A. Prescott, William W. Lytton.

**Keywords:** pain, spinal cord, spinal dorsal horn, computer modeling, degeneracy

## Abstract

Pain-related sensory input is processed in the spinal dorsal horn (SDH) before being relayed to the brain. That processing profoundly influences whether stimuli are correctly or incorrectly perceived as painful. Significant advances have been made in identifying the types of excitatory and inhibitory neurons that comprise the SDH, and there is some information about how neuron types are connected, but it remains unclear how the overall circuit processes sensory input or how that processing is disrupted under chronic pain conditions. To explore SDH function, we developed a computational model of the circuit that is tightly constrained by experimental data. Our model comprises conductance-based neuron models that reproduce the characteristic firing patterns of spinal neurons. Excitatory and inhibitory neuron populations, defined by their expression of genetic markers, spiking pattern, or morphology, were synaptically connected according to available qualitative data. Using a genetic algorithm, synaptic weights were tuned to reproduce projection neuron firing rates (model output) based on primary afferent firing rates (model input) across a range of mechanical stimulus intensities. Disparate synaptic weight combinations could produce equivalent circuit function, revealing degeneracy that may underlie heterogeneous responses of different circuits to perturbations or pathological insults. To validate our model, we verified that it responded to reduction of inhibition (i.e. disinhibition) and ablation of specific neuron types in a manner consistent with experiments. Thus validated, our model offers a valuable resource for interpreting experimental results and testing hypotheses *in silico* to plan experiments for examining normal and pathological SDH circuit function.

**Significance Statement:** We developed a multiscale computer model of the posterior part of spinal cord gray matter (spinal dorsal horn), involved in perception of touch and pain. The model reproduces several experimental observations and makes predictions about how specific types of spinal neurons and synapses influence projection neurons that send information to the brain. Misfiring of these projection neurons can produce anomalous sensations associated with chronic pain. Our computer model will not only assist in planning future experiments, but will also be useful for developing new pharmacotherapy for chronic pain disorders, connecting the effect of drugs acting at the molecular scale with emergent properties of neurons and circuits that shape the pain experience.

## Introduction

The spinal dorsal horn (SDH) plays a crucial role in processing somatosensory information. Sensory input conveyed to the SDH via primary afferents is processed by local excitatory and inhibitory interneurons before being relayed to the brain (Todd, 2010). Superficial layers of the SDH, laminae I-II, receive noxious input responsible for nociceptive pain. Deeper layers receive predominantly innocuous, low-threshold tactile input, but that input can be mistakenly perceived as painful if it is misprocessed (Prescott et al., 2014), resulting in mechanical allodynia, which is a common symptom in chronic pain conditions (Jensen and Finnerup, 2014).

The classic *Gate Control Theory* emphasized the importance of pain processing at the spinal level, particularly gating achieved through synaptic inhibition (Melzack and Wall, 1965). It is now well established that inhibition is reduced following nerve injury (for review see Prescott, 2015). This so-called disinhibition arises through different mechanisms (Coull et al., 2003; Zeilhofer et al., 2012) and differentially modulates spiking in excitatory and inhibitory interneurons (Lee et al., 2019). Early efforts to classify excitatory and inhibitory neurons by their spiking pattern, morphology, and neurochemical phenotype (Grudt and Perl 2002; Prescott and De Koninck 2002; Ruscheweyh and Sandkühler 2002; Yasaka et al. 2010) have been augmented more recently by molecular techniques to identify and manipulate genetically-defined neuron types (Hantman et al., 2004; Duan et al., 2014; Petitjean et al., 2015, 2019; Abraira et al., 2017; Zhang et al., 2018; Peirs et al., 2020, 2021). However, genetically-defined neuron types are often heterogeneous and extra steps must be taken to subdivide them into functionally homogeneous groups (Gradwell et al., 2021; Peirs et al., 2021). Decades of research have revealed SDH circuitry to be much more complex than envisioned by Melzack and Wall (for reviews see Peirs and Seal, 2016; Lechner, 2017).

Despite this progress, many unknowns remain and making sense of available data is becoming increasingly difficult. Indeed, impressive advances in basic pain research have not yet translated into improved clinical outcomes (Yekkirala et al., 2017). Thus, new tools are required to bridge persistent knowledge gaps. One way to take fuller advantage of available data is through multiscale computational modeling (O’Leary et al., 2015; Lytton et al., 2017; Hunt et al., 2018). Compared with other areas of neuroscience, pain research has seen relatively scant computational investigation. Various aspects of SDH neuron and circuit function have been modeled (Britton and Skevington, 1989; Prescott et al., 2006; Le Franc and Le Masson, 2010; Arle et al., 2014; Zhang et al., 2014, 2015; Ratté et al., 2015; Balachandar and Prescott, 2018; Crodelle et al., 2019) but even the most sophisticated circuit models consider only a small fraction of available experimental data.

Here, we developed a circuit model of lamina I-III that carefully incorporates the latest experimental data. Our conductance-based neuron models reproduce the characteristic spiking patterns of SDH neurons. The model also accounts for the differential input each neuron type receives from low- and high-threshold primary afferents. Using a genetic algorithm, synaptic weights were tuned to reproduce experimentally-observed projection neuron responses to mechanical stimuli of various intensities. This parameter search strategy revealed that equivalent SDH circuit function can be achieved with disparate synaptic weight combinations (solutions), but only a subset of solutions satisfied additional criteria not applied during fitting. Our top model reproduced several experimental observations including quantitative effects of disinhibition on SDH output, and qualitative effects of ablating specific neuron types. Our results emphasize the roles of inhibitory and excitatory interneurons in gating low-threshold input to projection neurons, and predict that targeting these upstream, polysynaptic pathways will attenuate effects of disinhibition and reduce allodynia. Finally, the model makes testable predictions regarding synaptic- and cellular-level targets for combating pathological disinhibition. This publicly-available model provides a resource for testing hypotheses *in silico* by linking molecular-level changes with emergent properties at the cellular- and circuit-levels to disentangle multiscale mechanisms underlying clinical symptoms.

## Materials & Methods

### SIMULATION ENVIRONMENT

We developed a realistic SDH model that reproduced circuit output in rodent dorsal horn during mechanical stimulation under normal conditions, pathological conditions, and following experimental ablation of specific spinal neuron types. Simulations were conducted with NEURON 8.0 (Hines and Carnevale, 2001; Carnevale and Hines, 2006) using NetPyNE 0.9.7 (Dura-Bernal et al. 2019b), http://netpyne.org, with an integration step of 25 μs at temperature 37°C. Model building, parameter optimization, and some of the model analysis were also performed using NetPyNE 0.9.7. The network consisted of 409 neurons from 14 population subtypes. Five seconds of simulation time took 23.4 min to run on one processor with Intel 1.2 GHz Core m3. Parallel simulations were conducted with 80 cores. Approximately 750,000 network simulations and 2500 single neuron simulations were run to develop this model.

### MECHANICAL STIMULATION AND PRIMARY AFFERENT MODELS

The SDH model was fit to experimental responses of primary afferents and spinal projection neurons to mechanical stimulation of the mouse paw at 50, 100, and 200 mN (Murthy et al., 2018; Walcher et al., 2018; Allard, 2019). These forces in mN correspond to 5.1, 10.2, and 20.2 g. The withdrawal threshold in mice is ~4–6 g based on electronic von Frey stimulation (Deuis et al., 2017), which was the convention used for experimental data (Murthy et al., 2018; Walcher et al., 2018); withdrawal threshold using conventional von Frey testing is about 6x smaller (Deuis et al., 2017). Throughout our subsequent modeling work, we simulate 5 to 200 mN mechanical force where allodynia corresponds to responsiveness to ~20 mN (i.e. 2.0 g or 0.3 g with electronic or conventional von Frey testing, respectively).

Responses of primary afferents, including Aβ, Aδ, and C-fibers, to mechanical stimulation were not simulated using conductance-based models, but were instead treated as random (i.e. Poisson) input spike trains whose rates, which vary dynamically during the 5 s-long stimulus, depend on mechanical stimulus intensity. The average rate and temporal profile were matched to *in vivo* recordings from mice (Murthy et al., 2018; Walcher et al., 2018). Specifically, we used *polyfit* (NumPy, Python) to reproduce the dynamic firing rate of Aβ fibers in response to five second mechanical stimulation at 100 mN (Walcher et al. 2018), as well as the dynamic rates of Aδ- and C-fibers in response to five second mechanical stimulation at 400 mN (Murthy et al. 2018). This response was then scaled to produce different firing rates at different stimulus intensities (**Fig. 1A, top**). Specifically, for C and Aδ fibers, the response at 400 mN was scaled by 0.001, 0.125, 0.25, and 0.5 to produce appropriate rates for <50, 50, 100 and 200 mN, respectively. According to Walcher et al. (2018), mean firing rates of Aβ-fibers plateau at around 60 mN (indicating their preference for low-intensity stimuli), so the dynamic firing rate of model Aβ-fibers at 50, 100, and 200 mN were all scaled by 1. During low-threshold stimulation (<50 mN), Aβ fibers were scaled linearly from 0.1 to 1 for 5 to 50 mN, respectively. Finally, using the dynamic firing rates of each fiber type, the spike trains for each primary afferent neuron (Aβ: n=20, Aδ: n=20, C: n=160) was created for each stimulus intensity using an inhomogeneous Poisson process. The bottom panel of **Figure 1A** shows the average firing rate for each primary afferent population. C fibers were subdivided into peptidergic (C,TRPV1; n=80) and non-peptidergic (C,IB4; n=80) subpopulations because only peptidergic C fibers directly connect to spinal projection neurons (Peirs and Seal, 2016; Lechner, 2017).

**Figure 1.**
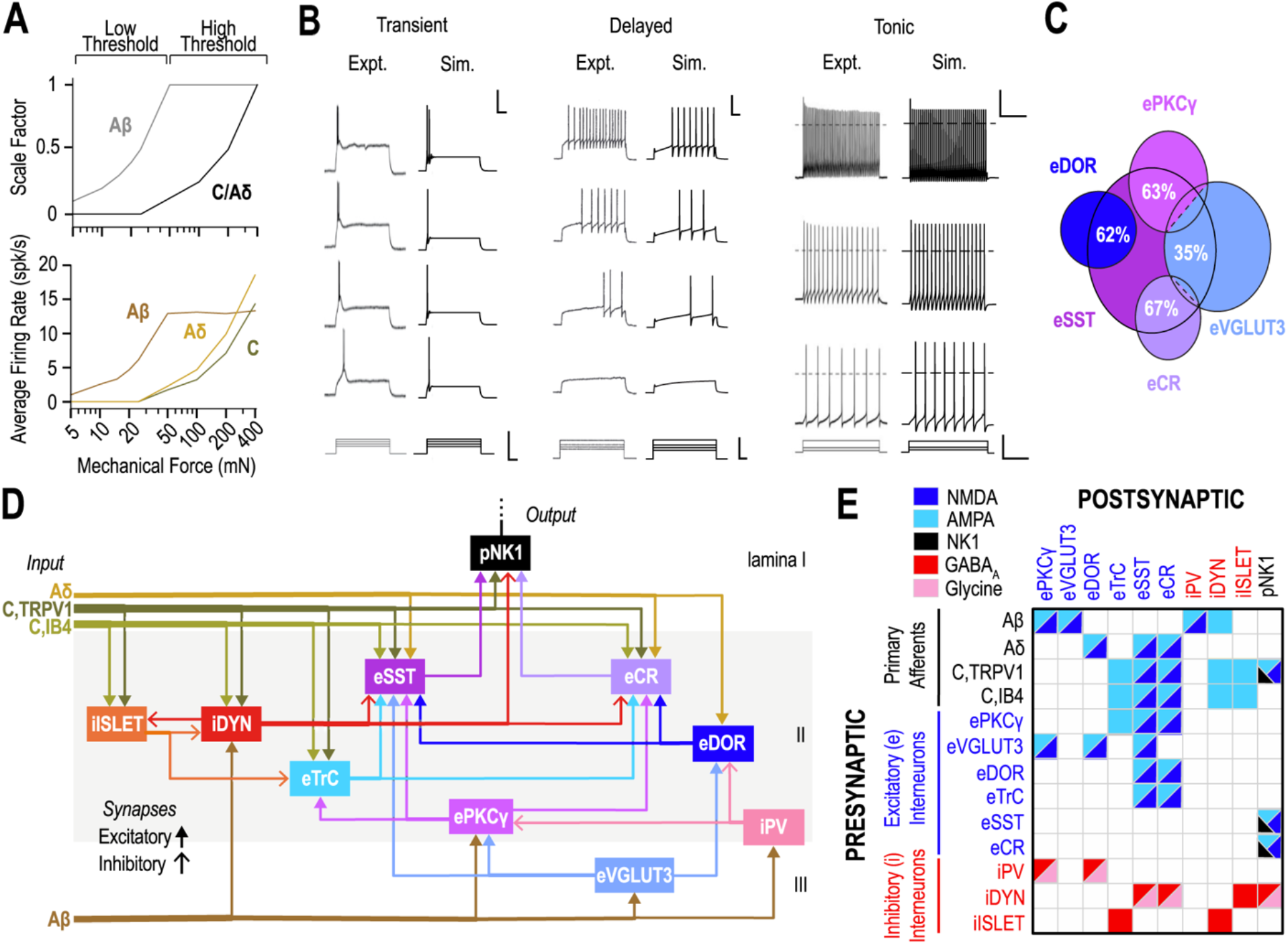
SDH circuit model. **(A)** Scale factor (top) and average firing rates (bottom) of model primary afferents (Aβ, Aδ, and C fibers) in response to mechanical stimulation. **(B)** Firing patterns of transient firing eTrC neurons (left), delayed firing ePKC*γ*, eVGLUT3, eDOR, eSST, and eCR neurons (center) and tonic firing iPV, iDYN, and iISLET neurons (right). Dashed line on the right panel indicates 0 mV. Scale bars: 150 ms, 40 mV (top) or 200 pA (bottom). **(C)** Gene expression overlap among excitatory interneurons. The percentage of overlap with eSST (white) is based on (Cheng et al., 2017) for eVGLUT3, (Gutierrez-Mecinas et al., 2016) for ePKC*γ* and eCR, and (Wang et al., 2018b) for eDOR. Overlap of eVGLUT3 with ePKC*γ*/eCR is based on (Peirs et al., 2015). **(D)** SDH circuit diagram. Primary afferent inputs (earth shades) include low-threshold afferents (Aβ) and high-threshold afferents (Aδ, peptidergic C (TRPV1), and non-peptidergic C (IB4)), which connect to excitatory interneurons *(e,* blue shades), inhibitory interneurons *(i,* red shades) and projection neurons *(p,* black) labeled by their gene expression or other defining property: CR, calretinin; DOR, delta opioid receptor; IB4, isolectin B4; NK1, neurokinin-1; PKC*γ*, protein kinase C gamma; PV, parvalbumin; SST, somatostatin; DYN, dynorphin; TrC, transient central; TRPV1, transient receptor potential V member 1; VGLUT3, vesicular glutamate transporter 3). Lamina are indicated on the right. **(E)** Synaptic connectivity matrix. Many synapses involve multiple receptor types, and are depicted accordingly.

### SPINAL CELL MODELS

The SDH contains excitatory and inhibitory interneurons whose axons remain in the spinal cord, as well as projection neurons whose axons extend to the brain. In the rodent spinal cord, experimental findings generally agree that delayed and transient firing are the most common patterns seen in excitatory interneurons, whereas tonic firing is most commonly observed in inhibitory interneurons (Punnakkal et al. 2014; Kardon et al. 2014; Lu and Perl 2005; Peirs et al. 2021; Todd 2017; Cathenaut et al. 2021; Cui et al. 2011; Duan et al. 2014; Yasaka et al. 2010; Wang et al. 2018; Smith et al. 2019). Therefore, starting from the morphology and conductance densities of spinal neurons used by Zhang et al. (2014), we reproduced delayed and transient firing in our excitatory neuron models and tonic firing in inhibitory spinal neurons (**Fig. 1B**). Each spinal neuron model consisted of three interconnected, single, cylindrical compartments representing the soma, dendrites, and axon initial segment (AIS). The morphology of excitatory and projection neurons were of similar proportions, which differed slightly from inhibitory neurons (see **Table 1**). Specifically, the soma compartment was larger for excitatory and projection neurons (20 μm in diameter) than inhibitory neurons (10 μm), but the dendrites of inhibitory neurons were longer based on previous models of tonic firing spinal neurons by Melnick et al. (2004). Next, an A-type potassium current (K_A_), which activates at subthreshold membrane potentials and delays spike onset, was added to excitatory neurons. Conductance density parameters in excitatory neurons (see **Table 2**) were then optimized to reproduce the spiking pattern, including the spike count and delay to first spike, across a range of current injection intensities applied to the soma through an IClamp point process. Transient central excitatory neurons (eTrC) produce 1–2 spikes within 100 ms of current onset (**Fig. 1B**, left). In delayed firing neurons, the latency to first spike decreases with increasing current (**Fig. 1B**, center). Spike height in these neurons was 107 mV from the starting voltage, similar to the experimental value of 110 mV. Tonic firing inhibitory neurons were simulated based on the morphology and conductance densities of previous spinal neuron models (Melnick et al., 2004; Zhang et al., 2014) and thus, exhibited a similar tonic firing profile to experimental data (**Fig. 1B,** right). Spinal projection neurons were also based on previous models (Aguiar et al., 2010; Zhang et al., 2014). Membrane capacitance for all spinal neuron models was 1.0 μF/cm^2^. Conductance densities are summarized in **Table 2.**

**Table 1.** Morphology of excitatory, inhibitory, and projection neuron models.

| Cell type | Section | Length<br>($\mu\text{m}$) | Diameter<br>($\mu\text{m}$) | Axial resistance<br>(ohm*cm) |
| --- | --- | --- | --- | --- |
| Excitatory interneurons (ePKCγ,<br>eVGLUT3, eTrC, eDOR, eSST,<br>eCR) | Dendrite | 400 | 3 | 150 |
|  | Soma | 20 | 20 | 150 |
|  | AIS | 9 | 1.5 | 150 |
| Inhibitory interneurons (iPV,<br>iDyn, iSLET) | Dendrite | 1371 | 1.4 | 80 |
|  | Soma | 10 | 10 | 80 |
|  | AIS | 30 | 0.74 | 80 |
| Projection neurons (pNK1) | Dendrite | 350 | 2.5 | 150 |
|  | Soma | 20 | 20 | 150 |
|  | AIS | 9 | 1.5 | 150 |

AIS, axon initial segment.

**Table 2.** Ion channels in excitatory, inhibitory, and projection neuron models.

| Spinal neuron population | Ion channel | Conductance at dendrite (mS/cm<sup>2</sup>) | Conductance at soma (mS/cm<sup>2</sup>) | Conductance at AIS (mS/cm<sup>2</sup>) |
| --- | --- | --- | --- | --- |
| Delayed firing excitatory interneurons (ePKCγ, eVGLUT3, eDOR, eSST, eCR) | HH-type Na<sup>+</sup>-1 | 0.0 | 85.48 | 23.75 |
|  | HH-type Na<sup>+</sup>-2 | 0.0 | 0.1652 | 30 |
|  | K<sub>dr</sub> | 9.60x10<sup>-3</sup> | 0.111 | 15.47 |
|  | HH-type K<sup>+</sup> | 144 | 4.3 | 304 |
|  | K<sub>A</sub> | 0.933 | 10.9 | 112 |
|  | K<sub>Ca</sub> | 2 | 2 | 0.0 |
|  | Leak | 9.6x10<sup>-4</sup> | 9.6x10<sup>-4</sup> | 9.6x10<sup>-4</sup> |
| Transient firing excitatory interneurons (eTrC) | HH-type Na<sup>+</sup>-1 | 0.0 | 0.0 | 0.0 |
|  | HH-type Na<sup>+</sup>-2 | 0.0 | 306.6 | 5147 |
|  | K<sub>dr</sub> | 206.1 | 0.0106 | 217.1 |
|  | K<sub>A</sub> | 0.1583 | 49.57 | 0.5 |
|  | K<sub>Ca</sub> | 2 | 2 | 0.0 |
|  | Leak | 4.2x10<sup>-2</sup> | 4.2x10<sup>-2</sup> | 4.2x10<sup>-2</sup> |
| Tonic firing inhibitory interneurons (iPV, iDYN and iSLET) | HH-type Na<sup>+</sup>-1 | 0.0 | 8 | 3450 |
|  | HH-type Na<sup>+</sup>-2 | 0.0 | 3450 | 8 |
|  | K<sub>dr</sub> | 34 | 4.3 | 76 |
|  | Leak | 1.1x10<sup>-2</sup> | 1.1x10<sup>-2</sup> | 1.1x10<sup>-2</sup> |
| Projection neurons (pNK1) | HH-type Na<sup>+</sup>-1 | 0.0 | 0.0 | 3450 |
|  | K<sub>dr</sub> | 36 | 1.075 | 76 |
|  | CaAN | 9.1x10<sup>-2</sup> | 0.0 | 0.0 |
|  | CaL | 3.0x10<sup>-2</sup> | 0.1 | 0.0 |
|  | K<sub>Ca</sub> | 1 | 0.1 | 0.0 |
|  | NaP | 0.0 | 0.1 | 0.0 |
|  | Leak | 4.2x10<sup>-2</sup> | 4.2x10<sup>-2</sup> | 4.2x10<sup>-2</sup> |

AIS, axon initial segment. HH, Hodgkin Huxley-type channels. K<sub>dr</sub>, delayed rectifier potassium channel. CaAN, slow calcium dependent cation channel. CaL, L-type high-threshold calcium channel. K<sub>Ca</sub>, slow calcium-dependent potassium channel. NaP, persistent sodium channel; K<sub>A</sub>, A-type potassium channel.

### SPINAL NEURON POPULATIONS

Inhibitory spinal interneurons (designated by “i”) were divided into three subtypes defined by their expression of parvalbumin (iPV) or dynorphin (iDYN), or by their islet morphology (iISLET). Some iPV neurons have islet morphology (Boyle et al., 2019), but in the model these two populations are implemented separately. Excitatory interneurons (designated by “e”) were subdivided into six subtypes defined by their expression of protein kinase C gamma (ePKC*γ*), vesicular glutamate transporter 3 (eVGLUT3), delta opioid receptor (eDOR), somatostatin (eSST), or calretinin (eCR) or by their transient-spiking pattern (eTrC). Previous experiments have shown substantial overlap in gene expression between different excitatory subtypes (**Fig. 1C**). Specifically, these studies have highlighted the overlap between different excitatory neuron populations expressing SST: 35% of VGLUT3-expressing neurons (Cheng et al., 2017), 63% of PKC*γ*-expressing neurons (Gutierrez-Mecinas et al., 2016), 67% of CR-expressing neurons (Gutierrez-Mecinas et al., 2016), and 62% of DOR-expressing neurons (Wang et al. 2018) have been shown to also express SST. Furthermore, (Peirs et al., 2015) reported a 25% and 7% overlap between VGLUT3-expression with CR or PKC*γ*, respectively, with little to no overlap between CR and PKC*γ* expression. In the SDH model, each of these neuronal subtypes are described as their own population, such that the eSST population is the subset of neurons which do not also express VGLUT3, PKC*γ*, CR, or DOR. Some eSST and eCR neurons have vertical morphology with dorsoventral and rostrocaudal dendrites (Huang et al., 2018; Gutierrez-Mecinas et al., 2019). Projection (p) neurons are defined by the expression of the neurokinin-1 (NK1) receptor (Todd et al., 2000) and are henceforth referred to as pNK1 neurons. Choi et al. (2021) recently identified subtypes of projection neurons that do not express the NK1 receptor, but Browne et al. (2019) found that 83% of neurons projecting to the parabrachial nucleus were NK1 positive.

The model contains inputs from 200 primary afferent fibers (20 Aβ, 20 Aδ, 80 C,IB4, and 80 C,TRPV1) and 209 spinal dorsal horn neurons (30 ePKC*γ*, 4 eVGLUT3, 30 eDOR, 10 eTrC, 15 eSST, 20 eCR, 15 iPV, 60 iDYN, 15 iISLET and 10 pNK1). The numbers of primary afferents and spinal neurons were approximated from experimental data. Specifically, the number of eSST, ePKC*γ*, eCR, eVGLUT3, iDYN and iPV neurons were assumed based on previous immunohistochemical expression data (Häring et al., 2018). The number of eDOR neurons was assumed to be the same as ePKC*γ* neurons based on additional immunohistochemical data (Wang et al., 2018). The number of eTrC neurons was assumed to be only a portion of the number of PKC*γ* and CR neurons because eTrC neurons are a subpopulation of PKC*γ* and CR neurons (Peirs et al., 2020). We set the number of iISLET neurons to be the same as iPV neurons. The proportion of excitatory (~55%) to inhibitory (~45%) spinal interneurons in our model is consistent with previous experimental work (Polgár et al., 2003, 2013). The number of projection (pNK1) neurons was assumed to be ~5% of the total number of spinal neurons based on quantitative experimental data showing that the majority of spinal neurons in lamina I-III are interneurons (Cameron et al., 2015; Todd, 2017). The total number of primary afferent fibers (~1000 fibers) was based on the rodent experimental data (Peyronnard et al., 1986; Le Bars et al., 2001) and the proportion of Aβ, Aδ, and C fibers (10%, 10%, 80%) was derived from experimental results (Le Bars et al., 2001). Similar to Crodelle et al. (2019), the number of neurons and afferent fibers in the SDH model represent one-fifth of the true number to reduce the computing load.

### SYNAPTIC CONNECTIVITY

Synaptic connectivity of the SDH model (**Fig. 1D,E**) was based on recent reviews of experimental data (Peirs and Seal, 2016; Lechner, 2017), with additional connections and neuron populations based on more recent observations (Petitjean et al. 2019; Wang et al. 2018; Smith et al. 2019). Historically, SDH circuitry was dissected using techniques like paired recordings (Lu and Perl, 2003, 2005; Graham et al., 2007), but more recent connectivity data come from genetic labeling, optogenetic, and ablation studies. Many experiments involve silencing/ablating different genetically-defined neuron populations and monitoring changes in sensitivity to mechanical stimuli. Results suggest that different spinal microcircuits are involved in different types of mechanical allodynia (e.g. static vs dynamic), and by doing so, provide clues about connectivity. For example, according to previous reviews (Peirs and Seal, 2016; Lechner, 2017), eCR neurons were thought to be presynaptic to eSST neurons, but a parallel arrangement is suggested by recent ablation studies indicating that eCR neurons selectively contribute to static allodynia following nerve injury (Duan et al., 2014; Petitjean et al., 2019) whereas eSST neurons contribute to both static and dynamic allodynia (Duan et al., 2014), and since some eCR neurons connect directly to pNK1 neurons (Petitjean et al. 2019; Smith et al. 2019); therefore, the eCR population was incorporated separately from eSST. Since eDOR neurons are selectively implicated in static allodynia (Wang et al. 2018), they were connected to both eSST and eCR neurons. In contrast, eVGLUT3 neurons are selectively implicated in dynamic allodynia (Cheng et al., 2017), and so they were connected to eSST neurons but not to eCR neurons. Since pharmacological inhibition of ePKC*γ* neurons reduced both types of allodynia (Petitjean et al., 2015), ePKC*γ* neurons were connected to both eSST and eCR neurons. It was also demonstrated that Aδ fibers exhibit monosynaptic or polysynaptic connections with eDOR and eSST, but not ePKC*γ* neurons (Wang et al. 2018), and this is therefore reflected in our model circuitry.

Synaptic connectivity also plays a role in the spike train variability of neurons across a population. This variability originates from input from *different* primary afferents whose spike trains have an equivalent rate but different patterns (i.e. each afferent spike train is generated from a separate Poisson process; see above). For two populations that are synaptically connected (see **Fig. 1E**), cells within each population are connected probabilistically such that each postsynaptic cell in population Y receives input from a different (partially overlapping) subset of presynaptic cells in population X, and each presynaptic cell in population X provides output to a different (partially overlapping) subset of cells in population Y. Each neuron in the presynaptic population has a 20% probability of forming a connection with a neuron in the postsynaptic population. We assumed a 20% probability of connection based on the sparse connectivity revealed by patchclamp recordings, which demonstrated that only 10-20% of simultaneously recorded pairs were synaptically connected (Lu and Perl, 2003, 2005; Lu et al., 2013).

### SYNAPSE MODELS

Excitatory synaptic transmission was mediated by AMPA, NMDA, and NK1 receptors whereas inhibitory synaptic transmission was mediated by GABA_A_ and glycine receptors. Therefore, AMPA and NMDA receptors were implemented at each excitatory connection and GABA_A_ and glycine receptors were implemented at each inhibitory connection, with some exceptions: NMDA receptors were not included at C–eTrC, C–iDYN, and C–iISLET synapses; glycine receptors were not included in iISLET–eTrC, iDYN–iISLET, and iISLET–iDYN synapses (Lu and Perl, 2003). NK1 receptors were included exclusively at the C,TRPV1-pNK1, eSST-pNK1, and eCR-pNK1 synapses based on experimental findings showing overlapping expression of substance P with SST (Gutierrez-Mecinas et al., 2017) or CR (Gutierrez-Mecinas et al., 2019). The different types of synapses formed between each presynaptic and postsynaptic population are summarized in **Fig. 1E**. Synapses were modeled using the Exp2Syn mechanism, which is defined by rise and decay time constants, and scaled by a synaptic weight. Rise and decay time constants (ms) were: AMPA 0.1, 5; NMDA 2, 100; NK1 100, 1000; GABA_A_ 0.1, 20; glycine 0.1, 10. The inhibitory reversal potential (*E*_inh_) was −70 mV and excitatory reversal potential (*E_exc_*) was 0 mV. Synaptic weights for each presynaptic to postsynaptic connection were tuned as described below.

### MODEL OPTIMIZATION

Synaptic weights were tuned using a genetic algorithm (GA) similar to previous studies (Neymotin et al. 2017; Dura-Bernal et al. 2015; Dura-Bernal et al. 2019a). We used a high-performance computing (HPC) cluster (Google Cloud Platform, https://cloud.google.com/; OIST Deigo, https://groups.oist.jp/scs/deigo) to efficiently optimize the model through parallelization of the GA, or more specifically, of the model evaluations at each generation (Sivagnanam et al., 2020). Parallel implementation of the GA over a large HPC cluster was achieved through the NetPyNE tool, which is interfaced with a Python-based optimization package *(Inspyred;* https://pypi.org/project/inspyred) and an open-source HPC job scheduling system *(Slurm;* https://slurm.schedmd.com). Our GA used tournament selection, uniform crossover, Gaussian mutation, and generational replacement (Carlson et al., 2014; Dura-Bernal et al., 2017). The population size, maximum number of generations, and mutation rate were set to 150, 100, and 0.4 for all GA fitting. Synaptic weights (35 values in total) were chosen by minimizing an error function (fitness corresponds to the inverse of the error). The error function was defined as the difference between median experimental (*expt*) and model pNK1 firing rates evoked by mechanical stimulation at 50, 100 and 200 mN, and the lack of pNK1 firing at 10 and 25 mN, quantified as,

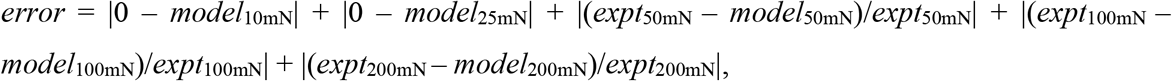

Synaptic weights for population initialization were randomly drawn from a beta distribution ranging from 1.0E-8 to 0.5. Weights for NK1 synapses were capped at 1.0E-6. The best solution, or set of 35 synaptic weights, was chosen based on the smallest error value after 150 generations (the GA stopping criterion). Median pNK1 firing rates had to fall within the interquartile range (IQR, 25th-75th percentile) of experimental data (IQR: 50 mN = (0.27–5.56) spk/s, 100 mN = (0.48–11.39) spk/s, 200 mN = (2.99–21.96) spk/s) and exhibit normal intensity discrimination (i.e. median pNK1 firing rate at 200 mN > 100 mN > 50 mN > 25 mN > 10 mN). If these convergence criteria were not met by 150 generations, the GA was stopped and run again with a different population initialization. Various iterations of the GA were run to fit our model to experimental data and to explore how synaptic weights changed for different population initializations. A subset of these solutions are shown in **Figure 3**. Further simulations were performed (see **Figs. 3, 4,** and **5**) to validate that our final solution of synaptic weights could reproduce other experimental data not used as part of the fitting process.

### STATISTICAL ANALYSIS

Data analysis was performed using NetPyNE 0.9.7 (Dura-Bernal et al. 2019b) and MATLAB R2021a (MathWorks). Five second-long SDH network responses were simulated. Similar to analysis from Allard (2019), individual data points from simulations represent the average firing rate of each simulated pNK1 neuron (n=10 per simulation run) calculated as the total number of spikes divided by the 5 second simulation period. Data with a non-Gaussian distribution are summarized as box plots showing the individual data points, and are summarized in the text as median (25th-75th percentile). Firing rate histograms (FRHs) were calculated using the kernel density estimation method (Shimazaki and Shinomoto, 2010). Briefly, spike trains across neurons of the same subpopulation were superimposed and the resulting spike sequence was convolved with a 100 ms-wide Gaussian kernel to obtain a smooth estimation of firing rate. Normally distributed data were compared using a t-test whereas a Mann-Whitney U test was used when data were not normally distributed. Pearson’s correlation coefficient *(r)* is stated for all linear regressions. The level of significance for all statistical tests was *p* < 0.05.

## Results

### Response of SDH model to mechanical stimulation

Having already set the parameters for single neuron models (see Methods), synaptic weights were tuned to reproduce pNK1 firing rates (circuit output) given afferent firing rates (circuit input) across a range of mechanical stimulus intensities (10, 25, 50, 100, and 200 mN). Based on the large number of synaptic weights to be tuned, a genetic algorithm was used to identify different sets of parameters (see Methods and **Fig. 3**). **Figure 2** shows sample responses from a circuit based on one set of synaptic weights. At 100 mN, activation of Aβ (brown), Aδ (gold), and C fibers (green) evoked spiking in all excitatory and inhibitory neuron populations except for iISLET neurons, and this resulted in pNK1 firing (black). Heterogeneity in the spike trains between neurons within a given population (**Fig. 2A**, raster plot) reflects the unique connectivity between each postsynaptic neuron and its presynaptic input (see Methods), as well as the randomness of spiking in primary afferents. There is no spontaneous activity in the SDH model and, as such, the model is quiescent prior to stimulation. At 50 mN, primarily Aβ fibers were activated (**Fig. 2B, left**), which was sufficient to activate some excitatory and inhibitory interneurons (blue and red hues, respectively), but caused only modest activation of projection neurons; ~40% of pNK1 neurons responded briefly, over a 0.5 s period, with 7-30 spikes. At 200 mN stimulation (**Fig. 2B, right**), activation of Aδ and C-fibers increased with no further increase in Aβ input. Because of these differences in afferent input firing rates (see **Fig. 1A**), deeper spinal neurons such as iPV and ePKC*γ* neurons, which receive predominantly Aβ input, showed minimal increases in firing as the stimulus intensity was increased from 50 to 200 mN. In contrast, neurons positioned more superficially, which receive direct input from Aδ and C fibers, showed substantial increases in firing between 50 to 200 mN.

**Figure 2.**
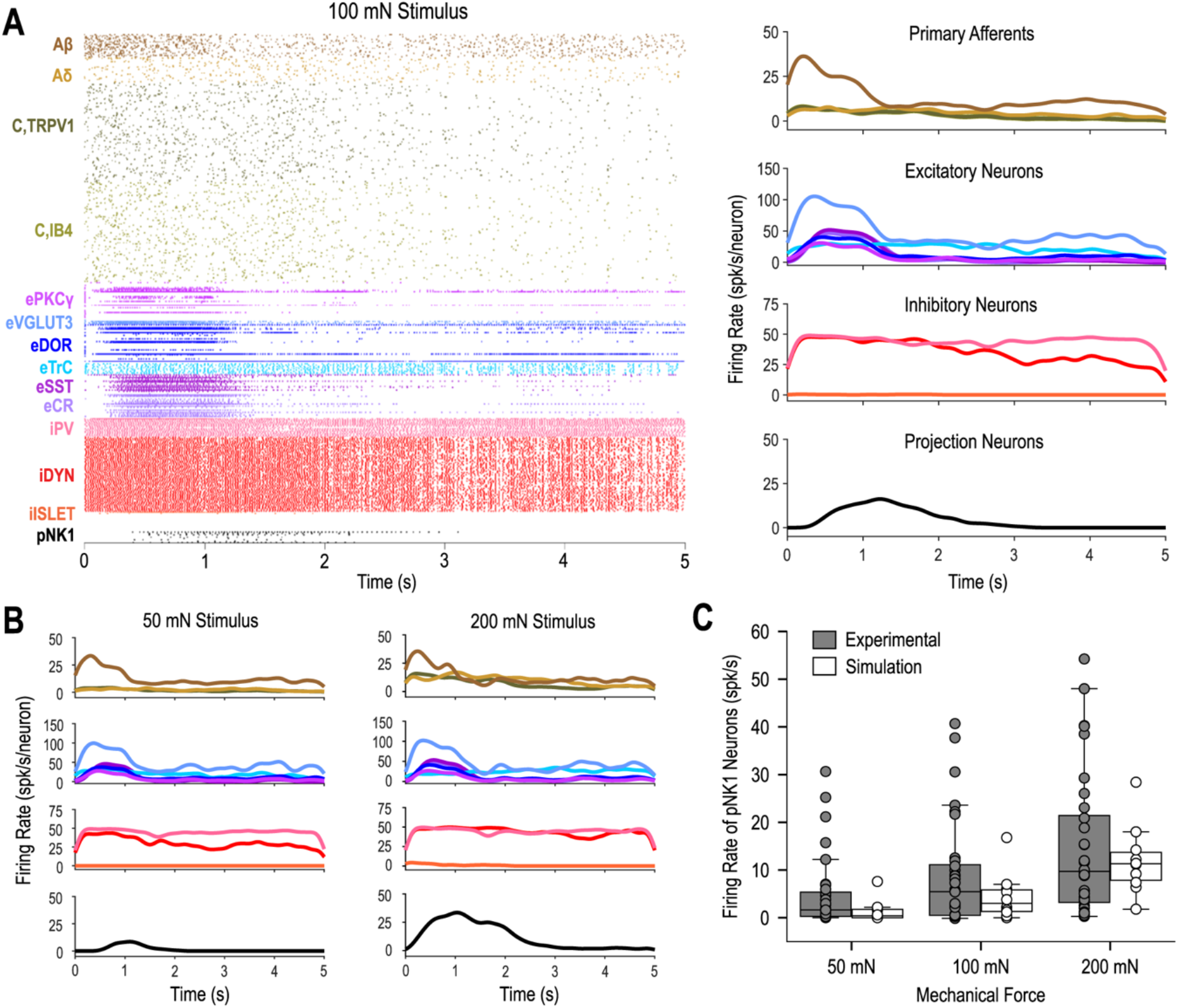
Sample responses of SDH model to mechanical stimulation. This corresponds to *circuit i* in Figure 3. **(A)** Representative response to 100 mN mechanical stimulation summarized as spike rasters for individual neurons (left) and firing rate histograms for each cell type (right). Each cell type is represented by the same color throughout the paper. Histograms were generated using a 100 ms-wide Gaussian kernel applied to the aggregate spike train for each cell type. **(B)** Firing rate histograms during 50 mN (left) and 200 mN (right) stimulation. **(C)** pNK1 firing rates (n=10) compared to experimental projection neuron firing rates (n=32 from Allard, 2019).

**Figure 2C** compares experimental firing rates from lamina I NK1 projection neurons during mechanical stimulation (Allard, 2019) with firing rates of model pNK1 neurons after optimizing synaptic weights. At each stimulus intensity tested, the model (white) reproduced the firing rates of real (gray) projection neurons (50 mN: experiment = 1.63 (0.27–5.56) spk/s, model = 0.40 (0–1.80) spk/s; 100 mN: experiment = 5.46 (0.48–11.39) spk/s, model = 3.00 (1.20–6) spk/s; 200 mN: experiment = 9.70 (2.99–21.96) spk/s, model = 11.30 (7.40–14.20) spk/s; experiment: n=32, model: n=10).

For the remainder of this study, we treat pNK1 firing rates as a proxy measure for pain. Notwithstanding obvious caveats (e.g. the impact of cortical processing on pain perception), this approximation is justified by several observations. Selective ablation of NK1 receptor positive projection neurons in lamina I attenuates allodynia and hyperalgesia in rodents (Mantyh et al., 1997; Nichols et al., 1999), and their selective activation evokes behaviors associated with stress and anxiety (Choi et al., 2021). This suggests that pNK1 neuron activation is necessary and sufficient to produce pain, which together with evidence that pNK1 firing rates correlate with pain intensity (Bester et al., 2000; Andrew and Craig, 2002; Allard, 2019), supports the use of pNK1 neuron activation is an *in silico* proxy for pain.

### Degeneracy in the SDH model

Previous experiments and simulations in other systems have shown that distinct sets of synaptic weights can produce equivalent circuit output (Prinz et al., 2004; Goaillard et al., 2009). This so-called degeneracy is important for maintaining robust circuit function (Marder, 2011; Tang et al., 2012; Marder et al., 2015; Goaillard and Marder, 2021) insofar as variations in one synaptic weight can be offset by compensatory variations in one or more other weights. It must be appreciated, however, that despite yielding equivalent circuit output under normal conditions, different synaptic weight combinations may produce very different circuit outputs following perturbation (Sakurai et al., 2014; Haddad and Marder, 2018).

Consistent with such degeneracy, multiple iterations of our genetic algorithm yielded 16 sets of synaptic weights (solutions) capable of reproducing pNK1 responses to mechanical stimulation under normal conditions. Weight matrices from three examples are shown in **Figure 3A**. **Figure 3B** shows their pNK1 neuron firing rates for 10 to 200 mN mechanical stimulation. Notably, our original error function provided little information about low-threshold (Aβ) input because pNK1 neurons do not normally respond to weak stimuli; in other words, the constraint on synaptic weights associated with Aβ input was weak. However, pNK1 neurons are known to respond to Aβ input when synaptic inhibition is blocked (see Introduction). Therefore, we tested how circuits based on different synaptic weights responded to removal of synaptic inhibition, simulated here by setting all GABA and glycine weights to 0. In the disinhibited state, all but one circuit responded with extremely high pNK1 firing rates (**Fig. 3C**); the exception, *circuit i,* responded with elevated pNK1 firing rates more closely approximating experimental observations (**Fig. 3C**, dashed line; Allard 2019, Keller et al., 2007). Notably, disinhibited responses to strong stimulation have not been quantified experimentally, and so it is unclear what firing rates to expect. Moreover, intrinsic cell mechanisms that would tend to limit the maximum sustainable firing rate, like slow sodium channel inactivation, were not included in our neuron models.

**Figure 3.**
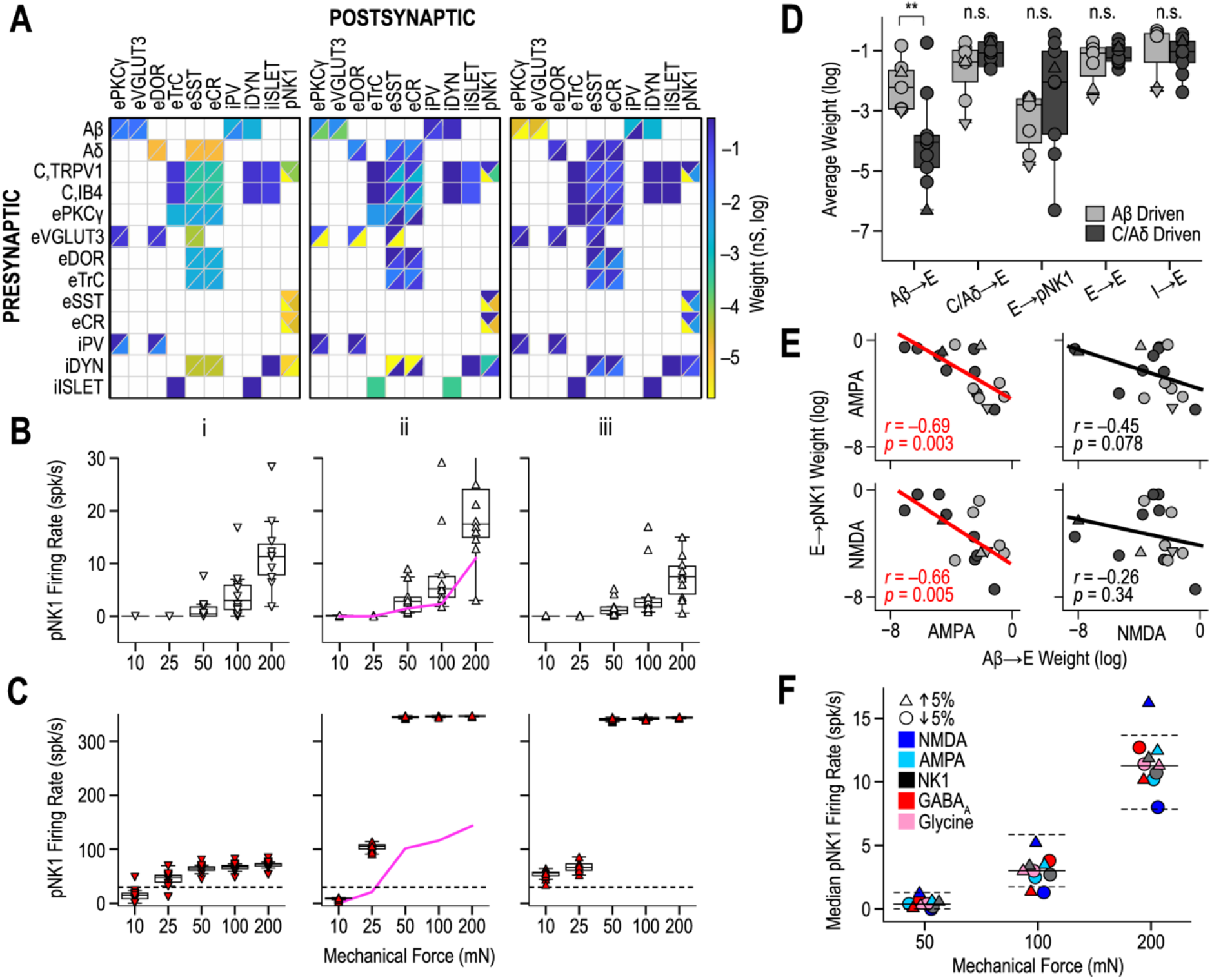
Degeneracy in the SDH circuit model. **(A)** Matrices of three synaptic weight combinations found by the GA fitting process. *(i)* SDH circuit reported in all other figures. *(ii)* Representative Aβ-driven circuit. (*iii*) Representative C/Aδ-driven circuit. **(B & C)** Response of models from A to 10-200 mN mechanical stimulation in normal conditions (B) and following complete blockade of GABA_A_ and glycine receptors (C). Dashed line in C represents experimentally measured median pNK1 firing rate in response to light touch following inhibitory receptor blockade (Allard, 2019). Thick pink lines shown for *circuit ii* in B and C report median pNK1 firing rate after reducing synaptic weights by a factor of 0.65 and 0.02 at deep and superficial synapses, respectively (see Results). **(D)** Average weight of different synaptic connections for circuits driven by Aβ (light gray, n=7) or C/Aδ (dark gray, n=9) fibers at low stimulus intensities. Light gray ▼, light gray ▲, and dark gray ▲ correspond to data from circuits i, ii, and iii in A, respectively. E, excitatory interneurons; I, inhibitory interneurons. **(E)** Correlations between AMPA and NMDA synaptic weights at Aβ→E vs E→pNK1 connections. Red linear regression lines indicate significant correlations (t-test). r, Pearson’s correlation coefficient. **(F)** Sensitivity analysis of *solution i.* Simulation was repeating after increasing (□) or decreasing (O) the strength of each type of synaptic weight by 5%. Solid and dashed lines show median and interquartile range, respectively, of pNK1 firing rates for original weights (same as in B*i*). Only three points fell outside the interquartile range, confirming that circuit function is relatively insensitive to small parameter changes.

Although *solution i* is considered our “best” outcome based on its disinhibited responses, analysis of the other solutions is nonetheless revealing. By removing Aβ inputs and re-testing the circuits, we determined if pNK1 neuron responses to weak (25 mN) stimulation in the disinhibited state were due to Aβ or C/Aδ input. In 7 circuits, Aβ input drove pNK1 neuron spiking whereas C/Aδ input was responsible in the other 9 circuits. Subdividing solutions based on this criterion, the synaptic weights between Aβ and excitatory interneurons was found to be significantly stronger in the former group than in the latter (**Fig. 3D;** *p*=0.018, t-test); other synaptic weights did not differ significantly between these groups. Further analysis revealed significant negative correlations between the strength of AMPA synapses between Aβ and excitatory interneurons and the strength of AMPA and NMDA synapses between excitatory interneurons and pNK1 neurons (**Fig. 3E**, left panels; Aβ→E(AMPA) vs. E→pNK1(AMPA): *r*=–0.69, *p*=0.003; Aβ→E(AMPA) vs. E→pNK1(NMDA): *r*=–0.66, *p*=0.005); in other words, the stronger the Aβ input to excitatory interneurons, the weaker the output of those interneurons to pNK1 neurons. This pattern is consistent with a variation in one parameter being offset by compensatory variation in other parameters, a hallmark of degenerate systems (see above). Notably, correlations between other synaptic weights that we considered were not significant (**Fig. 3E**, right panels; Aβ→E(NMDA) vs E→pNK1(AMPA): *r*=–0.45, *p*=0.078; Aβ→E(NMDA) vs. E→pNK1(NMDA): *r*=–0.26, *p*=0.34). These examples hint at the much deeper analysis that can be conducted on these sorts of data sets.

According to our results, many different sets of synaptic weights reproduced SDH output under normal conditions (i.e. the conditions to which the model was fit), but one set of synaptic weights, *solution i,* was superior in reproducing SDH output after disinhibition. Comparing solution *i* with other solutions is revealing. As evident in **Figures 3A** and **D**, synaptic weights tended to be weaker in *solution i* than in other solutions. In all solutions, synaptic excitation and inhibition were balanced, but in *solution i,* only weak synaptic inhibition was required to counterbalance weak synaptic excitation; therefore, when synaptic inhibition was removed, *circuit i* was less unbalanced (pNK1 firing rates increased less) than other circuits, reminiscent of the differential balance reported by (Lee et al., 2019). To test this, we shifted *solution ii* from strong excitation/strong inhibition to weak excitation/weak inhibition by reducing all synaptic weights except NK1 by a factor of 0.65 and 0.02 at deep and superficial synapses, respectively. Median pNK1 firing rates in this rescaled model are shown as solid pink lines on **Figure 3B** and **C**. Compared with the original *model ii,* pNK1 firing was significantly reduced in the disinhibited condition and more reasonably approximated experimental findings (Allard 2019), but was unchanged in the normal condition, as predicted. Since disinhibited responses were not included in the error function, and were not otherwise considered during fitting, testing the disinhibited responses serves to validate the model. Other solutions could be re-tuned to give reasonable disinhibited responses (see above), but *solution i* was considered our top candidate. Sensitivity analysis of *solution i* confirmed that a 5% increase or decrease in various synaptic weights caused only modest changes in pNK1 firing rates (**Fig. 3F**), thus excluding the possibility that circuit function relies on unrealistically precise parameter tuning. We therefore proceeded with *solution i* for all subsequent analysis.

### Effect of spinal neuron ablation

Having tentatively validated *solution i* on the basis of its disinhibited responses (see above), we proceeded to test how that model responded to other manipulations, namely ablation or silencing of specific types of neurons, for comparison with equivalent manipulations in experiments. For instance, Petitjean et al. (2015) demonstrated that ablation of PV neurons decreased mechanical threshold (i.e. increased sensitivity) to von Frey stimulation. Interestingly, mechanical allodynia in PV-ablated mice was reversed by pharmacological inhibition of PKC*γ* neurons. To examine the effects of iPV ablation in the SDH model, we set the number of iPV neurons to zero while leaving all other aspects of the model unchanged, and measured pNK1 neuron firing rates. Ablating iPV neurons increased median pNK1 firing rates from 0 to 39.1 (12.2-54.1) spk/s at 20 mN (**Fig. 4B*b***) and from 11.3 (7.4-14.2) to 70 (52.8-77.6) spk/s at 200 mN (**Fig. 4B*b’***). ePKC*γ* neurons were the excitatory interneurons most affected by the loss of iPV neurons (**Fig. 4*Ab,b’***). In agreement with experimental findings (Petitjean et al., 2015), removing ePKC*γ* neurons from our model mitigated the increased pNK1 firing caused by ablating iPV neurons (**Fig. 4*Bc,c*’**). This mitigating effect was especially strong for weak stimuli (20 mN) where pNK1 firing rates returned to normal (**Fig. 4B*c***). In the SDH model, iPV neurons inhibit ePKC*γ* and eDOR neurons, but not eSST neurons, which offer an alternate path for low-threshold inputs to reach pNK1 neurons (see **Fig. 1D**). Following iPV ablation, ablating eDOR neurons (**Fig. 4B*d,d’***) or eSST neurons (**Fig. 4B*e,e’***) also reduced the increased pNK1 firing, especially at 20 mN. Finally, simultaneously ablating ePKC*γ*, eDOR, and eSST neurons completely blocked the increase in pNK1 firing caused by iPV ablation at 20 mN (**Fig. 4*Bf***) and 200 mN (**Fig. 4B*f***). The input-output curve in **Figure 4C** summarizes the effect of different ablations on pNK1 firing at 0–200 mN and reveals that a ceiling effect is reached at 50 mN under certain conditions (e.g. following iPV ablation). In addition to confirming the importance of polysynaptic circuits for relaying low-threshold Aβ input to projection neurons, these simulations highlight that alternate routes are slightly different from each other (e.g. differently inhibited). By extension, these results predict that excitatory interneurons are an important target for combating allodynia, but that the efficacy of treatments targeting these interneurons may hinge on modulating all routes, lest Aβ input reach projection neurons via alternative, untargeted excitatory interneurons.

**Figure 4.**
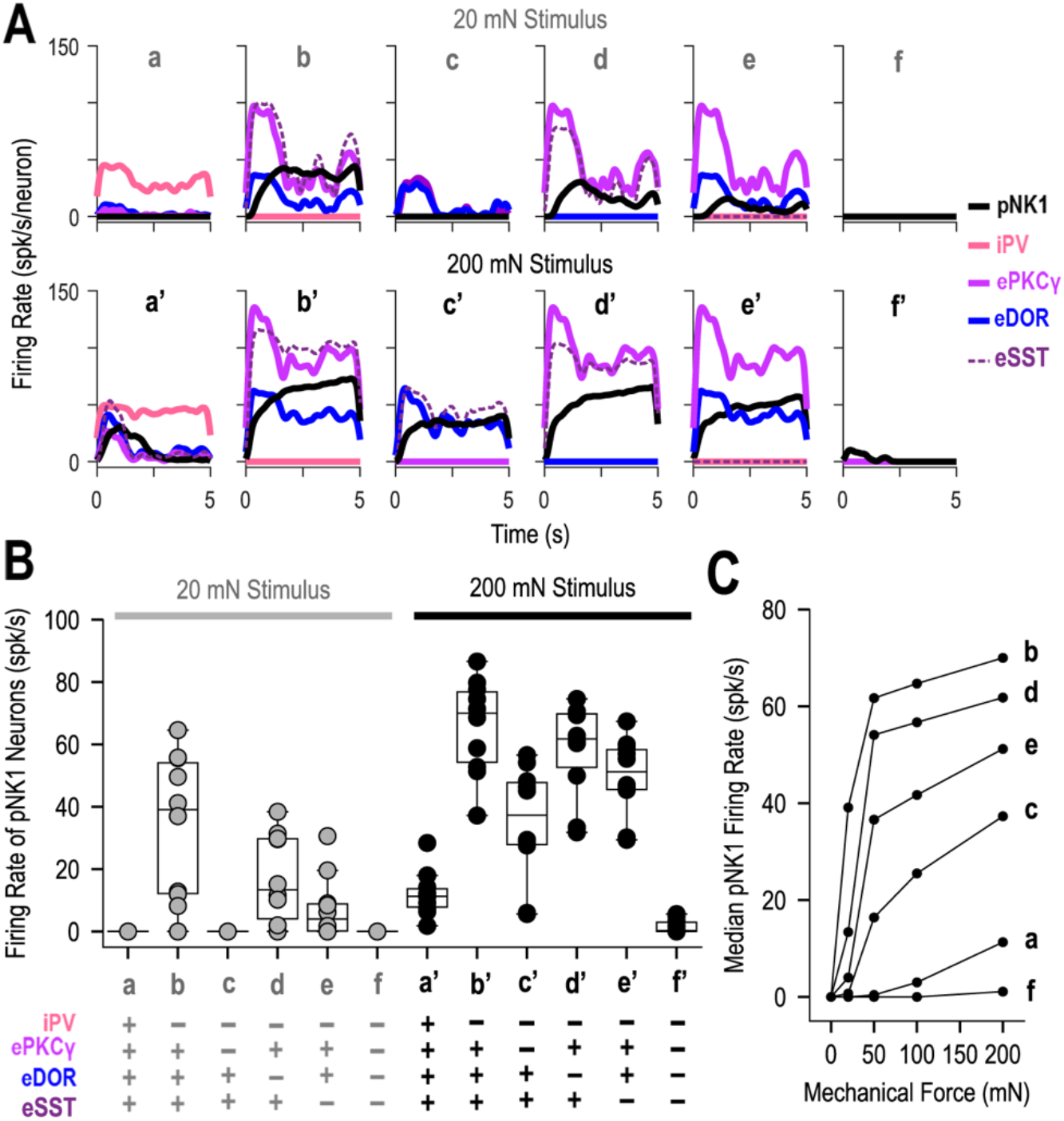
Effect of ablating iPV neurons on pNK1 firing depends on excitatory interneurons. **(A)** Firing rate histograms for response to 20 mN *(a-f)* and 200 mN (*a’-f’*) stimuli under control conditions (*a,a’*) and following ablation of iPV (*b,b’*) iPV and ePKC*γ* (*c,c’*), iPV and eDOR (*d,d’*), iPV and eSST (*e,e’*), or iPV, ePKC*γ*, eDOR and eSST (*f,f’*). **(B)** Summary of pNK1 firing rates under different ablation conditions. -, ablated; +, intact. **(C)** Input-output curves summarize median pNK1 firing rates from 0 to 200 mN across all conditions in B.

Next, we explored alternative routes and inhibitory mechanisms that control access of Aβ input to projection neurons. Selective ablation of iDYN neurons did not increase pNK1 firing at 20, 50, 100 or 200 mN (**Fig. 5A,** red), which is contrary to findings by Duan et al. (2014). We theorized that this stark difference from the effect of iPV ablation (**Fig. 5A,** pink; same as **Fig. 4**) was the result of SDH circuit degeneracy. Specifically, as both iPV and iDYN neurons inhibit major excitatory interneuron populations, we hypothesized that iPV- and iDYN-derived inhibition were functionally equivalent and could thus be traded-off without compromising circuit responses to mechanical stimulation. Indeed, iPV output weights were stronger than iDYN output weights in *solution i,* but this was not uniformly true of other solutions (**Fig. 3A**).

**Figure 5.**
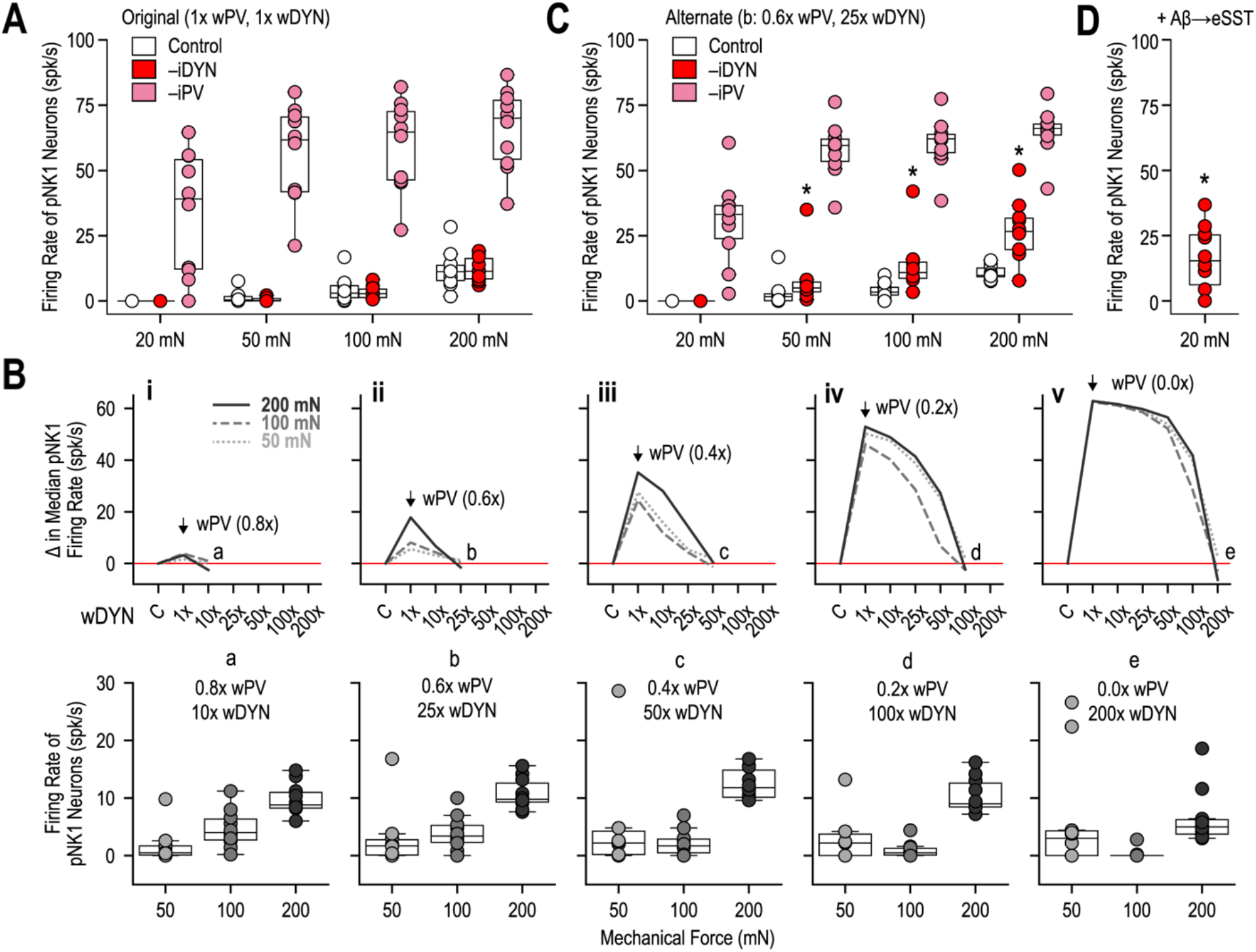
Trade-off between iPV- and iDYN-mediated inhibition. wPV, synaptic weights between iPV and ePKCγ/eDOR; wDYN, synaptic weights between iDYN and eCR/eSST. **(A)** Response of original SDH model (1x wPV, 1x wDYN; weights from Fig. 3A*i*) to 20, 50, 100, and 200 mN stimulation under control conditions (white) or following ablation of iDYN (red) or iPV (pink) neurons. **(B, top)** Increases in median pNK1 firing rates (corrected to control, *C)* following a 20% (*i*), 40% (*ii*), 60% *(iii),* 80% *(iv)* or 100% (*v*) reduction in wPV can be compensated for by concurrent increases in wDYN (x-axis). Full compensation of pNK1 firing rates is indicated by return of curves to the red line at points *a-e.* **(B, bottom)** Firing rates of pNK1 neurons at 50, 100, and 200 mN with compensated wPV/wDYN combinations. **(C)** The same simulation conditions as A but with alternate iPV/iDYN weight combination ***b*** (0.6x wPV, 25x wDYN). Ablating iDYN neurons had a significantly larger effect in alternate model (C) than in original model (A) for all stimulus intensities except 20 mN (*, p < 0.005, Mann-Whitney tests); effects do not differ significantly (p > 0.4) between A and C for control or iPV ablation conditions. **(D)** Response of pNK1 neurons to 20 mN following iDYN ablation and addition of direct Aβ to eSST input. * indicates significant (p < 0.005) increase in pNK1 firing rate compared to iDYN ablation at 20 mN in A and C.

To test our hypothesis, we systematically reduced the weights of iPV connections to ePKC or eDOR neurons (wPV), while increasing the weights of iDYN connections to eSST and eCR neurons (wDYN). Subtle decreases in wPV (a 20% reduction or 0.8x wPV) led to slight increases in pNK1 firing rates across all stimulus intensities which could be offset by a 10x increase in wDYN (**Fig. 5*Bi***, top). This weight combination of 0.8x wPV and 10x wDYN (***a***) resulted in pNK1 firing rates similar to control conditions (1x wPV, 1x wDYN) and this tuning maintained normal intensity discrimination, namely increased pNK1 firing with increasing stimulus intensity (**Fig. 5B*i***, bottom; pNK1 firing rates for ***a***: 50 mN = 0.5 (0.1–1.7) spk/s, 100 mN = 4.0 (2.7–6.4) spk/s, 200 mN = 8.8 (8.3–14.8) spk/s). A 40% decrease in wPV (or 0.6x wPV) led to greater increases in pNK1 firing rates that required a larger increase (25x) in wDYN to achieve full compensation (**Fig. 5B*ii***, top). Similar to ***a***, the weight combination of 0.6x wPV and 25x wDYN (***b***) resulted in pNK1 firing rates that were comparable to control conditions and maintained normal stimulus intensity discrimination (**Fig. 5B*ii***, bottom; pNK1 firing rates for ***b***: 50 mN = 1.7 (0.1–2.8) spk/s, 100 mN = 3.4 (2.3–5.3) spk/s, 200 mN = 9.8 (9.3–15.6) spk/s). Larger or complete decreases in wPV (0.4x, 0.2x, or 0x wPV) all caused progressively greater increases in median pNK1 firing rates and necessitated progressively larger increases in wDYN (50x, 100x, 200x, respectively) for full compensation (**Fig. 5B*iii-v***). However, with these weight combinations (***c**, **d***, and ***e***), compensated median pNK1 firing rates lost their stimulus intensity discrimination (**Fig. 2C**, gray). These results confirm our hypothesis that increased iDYN-mediated inhibition can offset decreased iPV-mediated inhibition to maintain normal SDH output, but only within certain limits.

Using weight combination ***b*** (0.6x wPV, 25x wDYN), our model demonstrated more substantial increases in pNK1 firing following iDYN ablation, especially for strong stimuli (≥ 50 mN), without compromising the effect of iPV ablation on pNK1 firing (**Fig. 5C**). These results suggest that iPV neurons are more important for allodynia, whereas iDYN are more important for hyperalgesia; however, such interpretations are limited by degeneracy of the SDH circuit (see **Fig. 3**). Previous experimental work by Duan et al. (2014) showed that iDYN ablation unmasked direct Aβ input to eSST neurons, presumably by removing presynaptic inhibition of Aβ fibers. Although we did not formally simulate presynaptic inhibition in our model, addition of a direct Aβ input to eSST neurons during iDYN ablation increased the response of pNK1 neurons at 20 mN (**Fig. 5D**), thus identifying a mechanism through which iDYN neurons may be implicated in allodynia.

In summary, the results of **Figure 5** demonstrate a trade-off between iPV- and iDYN-mediated inhibition that does not disrupt the functional integrity of the SDH circuit. This degeneracy was revealed by co-adjusting iPV and iDYN weights, and is reminiscent of the negative correlations presented in **Figure 3E**. Finally, ablations in **Figures 4** and **5** further validate the SDH model against experimental data, highlight the importance of different inhibitory and excitatory spinal interneuron populations for gating Aβ input to projection neurons, and predict that this gating of primary afferent input by spinal interneurons is especially important for processing low-threshold inputs.

### Effects of disinhibition

Several experimental studies have shown that blocking synaptic inhibition in the spinal cord can acutely reproduce the allodynia and hyperalgesia observed after nerve injury or in other neuropathic pain conditions (for review see Prescott, 2015). Furthermore, *in vivo* recordings have documented the associated increase in projection neuron firing after reducing synaptic inhibition (Keller et al., 2007; Allard, 2019). Effects of disinhibition on our SDH model were discussed briefly in **Figure 3**, but are fleshed out more fully here, taking into account the role of excitatory interneurons described above.

**Figure 6A** shows pNK1 firing rates before (white) and after partial (90%) and complete blockade GABA_A_ and glycine receptors (red and gray, respectively). Inhibitory receptor blockade (IRB) was simulated by setting all GABAergic and glycinergic synaptic weights to 10% or 0% of their original values. IRB caused increased pNK1 firing rates across all stimulus intensities, consistent with disinhibition causing hyperalgesia (increased pain in response to strong stimuli) and allodynia (pain in response to normally innocuous stimuli). Complete and partial IRB caused nearly equivalent pNK1 firing at high intensities (50-200 mN) whereas, at lower stimulus intensities, pNK1 firing rates were substantially higher after complete IRB than after partial IRB, which suggests that a small amount of residual inhibition can go a long way in attenuating SDH output (see below). The range of disinhibited pNK1 firing rates (0-85 spk/s) is consistent with experimental data (10-170 spk/s; Allard, 2019). Most notably, after IRB, innocuous stimulation (10-20 mN, depending on the degree of receptor blockade) evoked equivalent pNK1 firing rates as noxious stimulation (200 mN) under normal stimulus conditions, consistent with disinhibition causing mechanical allodynia.

**Figure 6.**
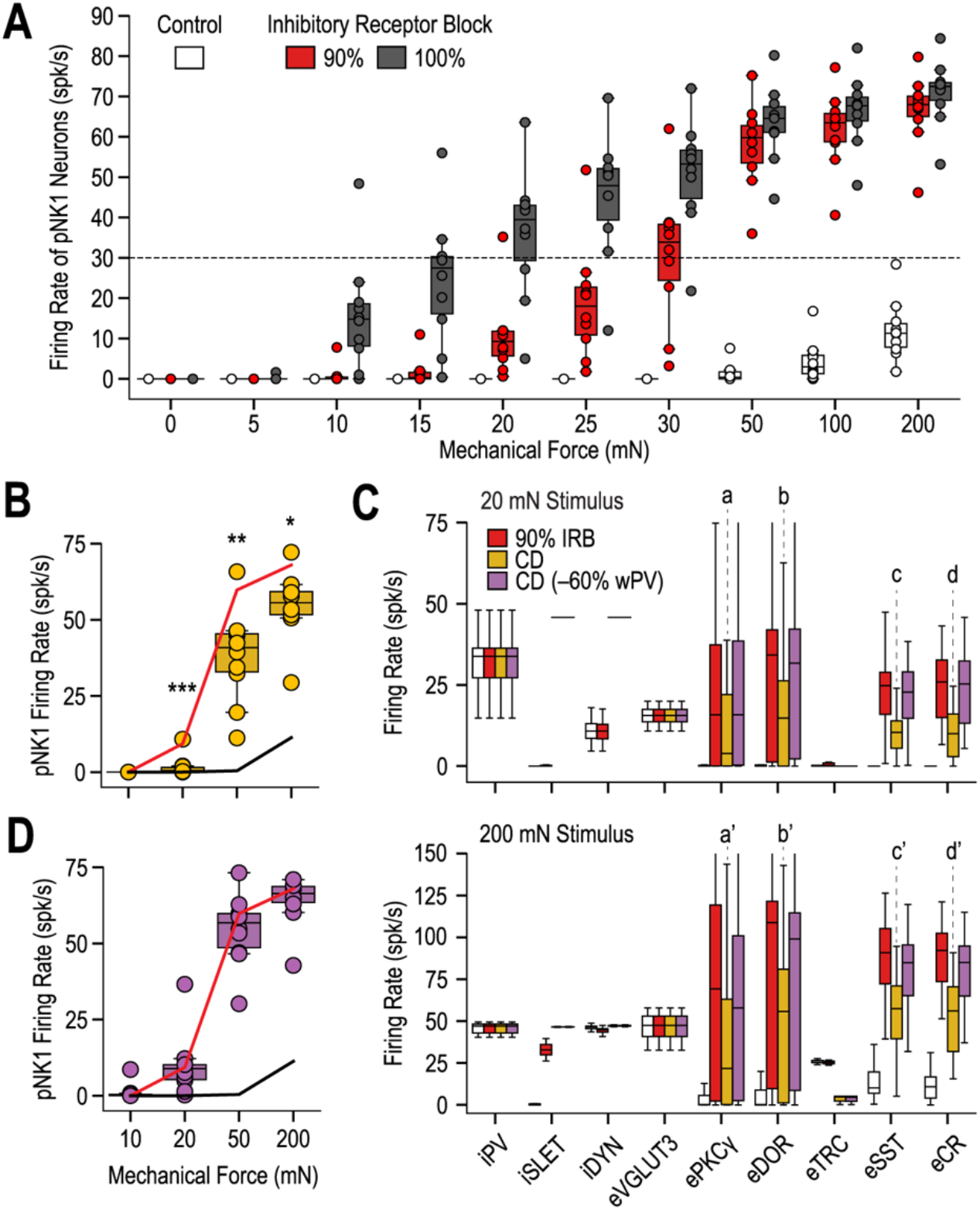
Response of SDH model to different forms of disinhibition. **(A)** Firing rates of pNK1 neurons (n=10) to 10-200 mN stimulation following partial (90%, red) or complete (100%, gray) inhibitory receptor blockade (IRB). The SDH model quantitatively reproduces experimental data after IRB (dashed line same as in Figure 3C). **(B)** Response of pNK1 neurons to 10, 20, 50, and 200 mN stimulation following a +25 mV shift in *E*_inh_ to simulate chloride dysregulation (CD). Lines show median pNK1 firing rates under control conditions (black) and after 90% IRB (red). Compared with 90% IRB, the increase in pNK1 firing due to CD was significantly less at 20 mN (***, p=0.0024), 50 mN (**, p=0.0073) and 200 mN (*, p=0.014; Mann-Whitney U tests). **(C)** Firing rates of different spinal interneuron populations following 90% IRB (red), CD (yellow), or CD and a 60% reduction in iPV weights (wPV) (purple). For 20 mN stimulation, increased firing due to CD was significantly less than for 90% IRB for eSST (c, p=0.0090) and eCR (d, p<0.001) neurons, but not quite for ePKC*γ* (a, p=0.18) or eDOR (b, p=0.10; Mann-Whitney U tests) neurons. Results were similar for 200 mN stimulation: ePKC*γ* (a’, p=0.083), eDOR (b’, p=0.039), eSST (c’, p=0.011), and eCR (d’, p<0.001). wPV was reduced in order to selectively increase ePKC*γ* and eDOR firing rates to values seen after 90% IRB and thereby test the consequences of altered excitatory input to downstream neurons. **(D)** Response of pNK1 neurons to CD plus a 60% reduction in wPV. Red and black lines are the same as in B. None of the responses deviated significantly from 90% IRB (p>0.4; Mann-Whitney tests). Overall, the results show that a small difference in inhibition leads to changes in excitatory input that compound at the circuit-level.

In neuropathic pain conditions, inhibition in the SDH is commonly compromised by chloride dysregulation (CD). CD stems from downregulation of the potassium-chloride cotransporter 2 (KCC2), which causes elevated intracellular chloride and a depolarizing shift in the anion reversal potential (which corresponds to *E*_inh_ in our model) (Coull et al., 2003). *In vivo* recordings have documented the effects of CD on different types of SDH neurons (Keller et al., 2007; Lee et al., 2019; Ferrini et al., 2020). IRB and CD, which can be induced artificially by blocking KCC2, appear to have similar effects on SDH interneurons (Lee and Prescott, 2015) but the impact on circuit-level function remains to be investigated. Using our model, we investigated the effect of shifting *E*_inh_ from −70 to −45 mV. A depolarizing shift in *E*_inh_ increased pNK1 firing rates during strong stimulation (50–200 mN), but caused more subtle increases in pNK1 firing during weak stimulation (20 mN) (**Fig. 6B**). The increase in pNK1 firing due to CD was smaller than that caused by 90% IRB. This finding is consistent with (Keller et al., 2007), who found that GABA_A_ receptor block led to slightly larger increases in projection neuron firing to innocuous touch than KCC2 blockade, but is inconsistent with other work showing that KCC2 blockade produces allodynia and that reversing CD reduces allodynia (Gagnon et al., 2013; Mapplebeck et al., 2019).

To investigate this further, we compared the firing rates of interneurons in the SDH circuit during weak (20 mN) and strong (200 mN) stimulation after disinhibition by 90% IRB or CD (red and gold, respectively on **Fig. 6C**). The most remarkable difference was that excitatory interneurons immediately presynaptic to pNK1 neurons (namely eSST and eCR neurons) fired much less after CD than after 90% IRB. Excitatory neurons further upstream (namely eTRC, ePKCγ and eDOR neurons) also fired less after CD than after 90% IRB, but the difference was smaller. This suggests that disinhibition has a compounding effect: disinhibiting upstream excitatory neurons leads to increased excitatory input to downstream neurons, which, if not balanced by inhibition, leads to progressive increases in excitatory input to each subsequent layer of the feedforward network. To test this explanation, we reduced the weight of iPV input ePKCγ and eDOR neurons by 60% (purple on **Fig. 6C**) to selectively increase ePKC*γ* and eDOR firing rates to values seen with global 90% IRB (red). As predicted, the elevated ePKCγ/eDOR output increased firing in downstream eSST and eCR neurons, which in turn increased pNK1 firing (**Fig. 6D**). Reminiscent of the sizable difference in pNK1 firing caused by a mere 10% difference in receptor blockade (see **Fig. 6A**), a small difference in disinhibition caused by CD and 90% IRB translated into a sizable difference in pNK1 firing via circuit-level effects. This may also help explain the greater impact of iPV-mediated inhibition compared with iDYN-mediated inhibition (see **Fig. 4**), notwithstanding effects of different synaptic weight combinations (see **Fig. 5**). It is, therefore, notable that most Aβ input reaches pNK1 neurons via at least two interposed neurons (Torsney and MacDermott, 2006). This compounding effect would be mitigated if inhibitory interneurons were also activated by excitatory interneurons, but they are activated exclusively by afferent input according to available evidence; however, absence of evidence is not evidence against, which is to say that more experiments are required to rule in/out excitatory interneuron connections to inhibitory interneurons.

### Mitigation of disinhibition by multiscale mechanisms

Given the importance of excitatory interneurons for relaying Aβ input to pNK1 neurons, we explored mechanisms through which the effects of disinhibiting those polysynaptic pathways could be mitigated. Capitalizing on our multiscale representation of the SDH, we first explored targets at the synaptic-level. NK1 receptor (NK1R) blockade has been shown to attenuate mechanical allodynia in rodents (Cahill and Coderre, 2002). As such, we simulated NK1R blockade by setting the synaptic weights at all NK1 synapses to zero (**Fig. 7**). When inhibition was intact, NK1R blockade had no impact on pNK1 responses to 20 mN stimulation (**Fig. 7A*b***) but it did reduce responses to 200 mN stimulation (**Fig. 7A*b’***). As previously shown, disinhibition unmasks significant Aβ input to pNK1 neurons (see **Figs. 4, 5,** and **6**). The increase in pNK1 firing caused by complete (100%) GABA_A_ and glycine receptor blockade (**Fig. 7A*c,c’***) was dramatically attenuated by NK1R blockade for 20 mN (**Fig. 7A*g***) and 200 mN (**Fig. 7A*g’***) stimulation. These findings are consistent with NK1R contributing to mechanical allodynia and hyperalgesia, respectively. Blockade of NK1 synapses between projection neurons and individual presynaptic connections revealed that NK1 synapses formed by eSST (**Fig. 7A*e/e’***) and eCR (**Fig. 7A*f/f’***) neurons onto pNK1 neurons were more vital for mitigation of disinhibition than those formed by C fibers (**Fig. 7A*d/d’***). Complete NK1R blockade was particularly effective in mitigating the effects of disinhibition at low stimulus intensities (20 mN), returning pNK1 firing to its baseline level. These findings are summarized in **Fig. 7B** and demonstrate our model’s prediction that NK1R blockade, a synaptic-level mechanism, can mitigate the effects of disinhibition.

**Figure 7.**
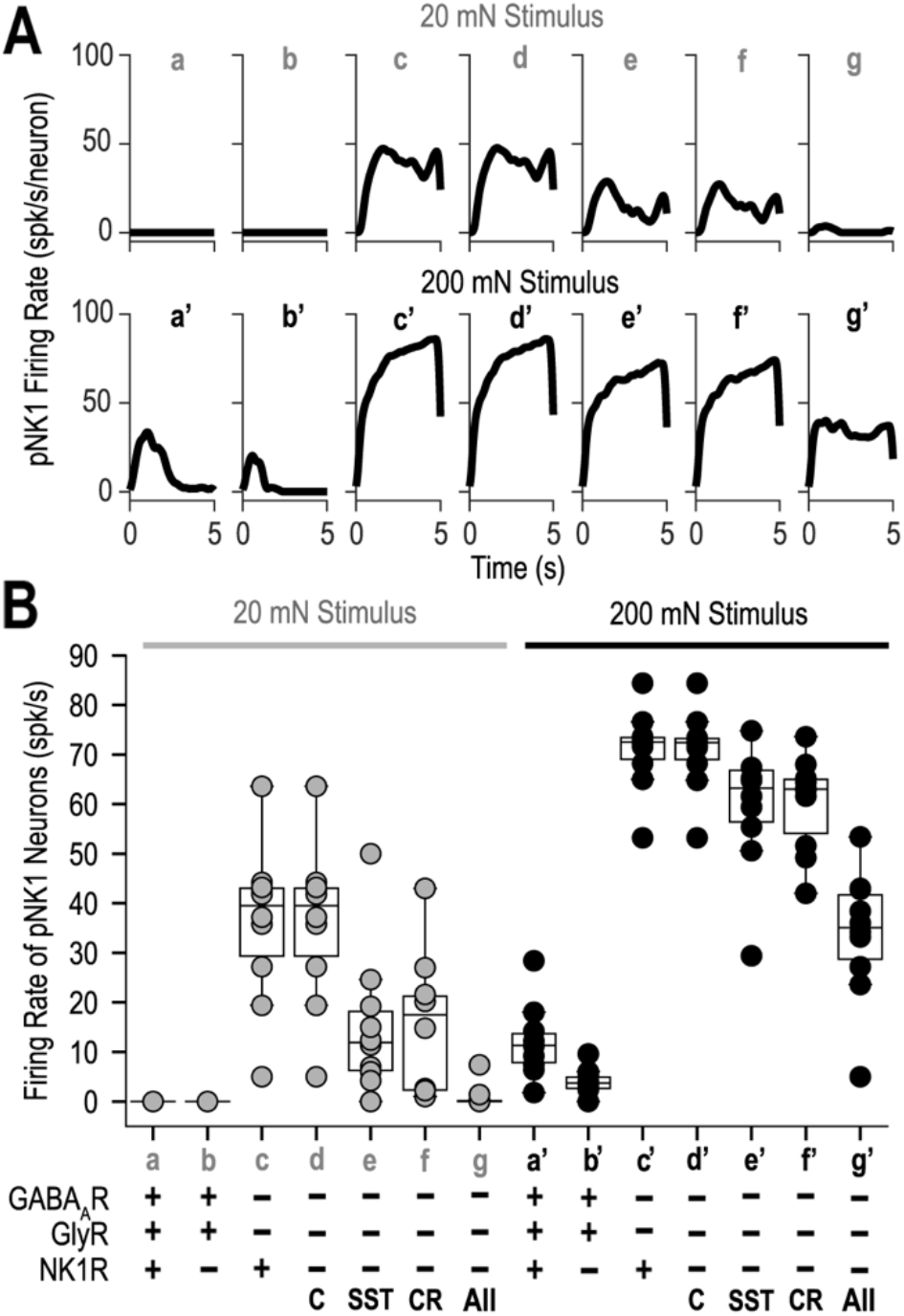
NK1 receptor blockade can mitigate effects of disinhibition on pNK1 firing. **(A)** Firing rates of pNK1 neurons under control conditions (20 mN: *a-d;* 200 mN: *a’-d’)* and after NK1 receptor blockade (*b/b’*), GABAA/glycine receptor blockade (*c/c’*), or simultaneous blockade of GABAA/glycine receptors and NK1 receptors specifically at C fiber *(d/d’*) eSST (*e/e’*), or eCR (*f/f’*) synapses onto pNK1 neurons. Simultaneous blockade of all NK1 synapses following disinhibition is shown in *g/g’*. **(B)** Summary of pNK1 firing rates under different simulation conditions. −, blocked; +, intact.

Whereas NK1R receptor blockade is predicted to reduce SDH output in response to noxious stimulation, the changes underlying allodynia could in principle be more selectively counteracted by targeting deep excitatory interneurons responsible for relaying Aβ input to pNK1 neurons (see **Fig. 4**), especially the ones furthest upstream (see **Fig. 6C**). Therefore, we explored the effects of reducing the excitability of excitatory interneurons. In the spinal dorsal horn, Kv4.2 and Kv4.3 subunits, which form A-type potassium (K_A_) channels, are selectively expressed by excitatory interneurons (Huang et al., 2005; Häring et al., 2018). K_A_ channels regulate neuronal excitability at subthreshold potentials and have been implicated in pain sensitivity (Chien et al., 2007; Duan et al., 2012; Zhang et al., 2018). We hypothesized that through control of intrinsic neuronal excitability, K_A_ channels represent a cellular-level (rather than synaptic-level) gating mechanism that may selectively modulate the allodynia produced by disinhibition.

Theoretically, reducing K_A_ channel conductance (gK_A_) should increase the firing rate of excitatory interneurons and thus lead to increased pNK1 firing. As predicted, reducing gK_A_ caused increased pNK1 firing at both 20 mN (**Fig. 8A**, top) and 200 mN (**Fig. 8A**, bottom). The median firing rate of pNK1 neurons increased from 0 to 82.2 spk/s at 20 mN and 11.30 to 95.90 spk/s at 200 mN following complete blockade of K_A_ channels (0% of baseline). The firing rates of pNK1 neurons following decreases in gK_A_ to 20% of its baseline at 20 mN or 40% of its baseline at 200 mN were comparable to that after inhibitory receptor blockade at the same respective stimulus intensities (**Fig. 6A**), suggesting that blockade of K_A_ channels could lead to both allodynia and hyperalgesia. Conversely, at 160% of baseline gK_A_, the median pNK1 firing rate at 200 mN decreased to 0.9 spk/s where it plateaued indicating a floor effect of K_A_ enhancement.

**Figure 8.**
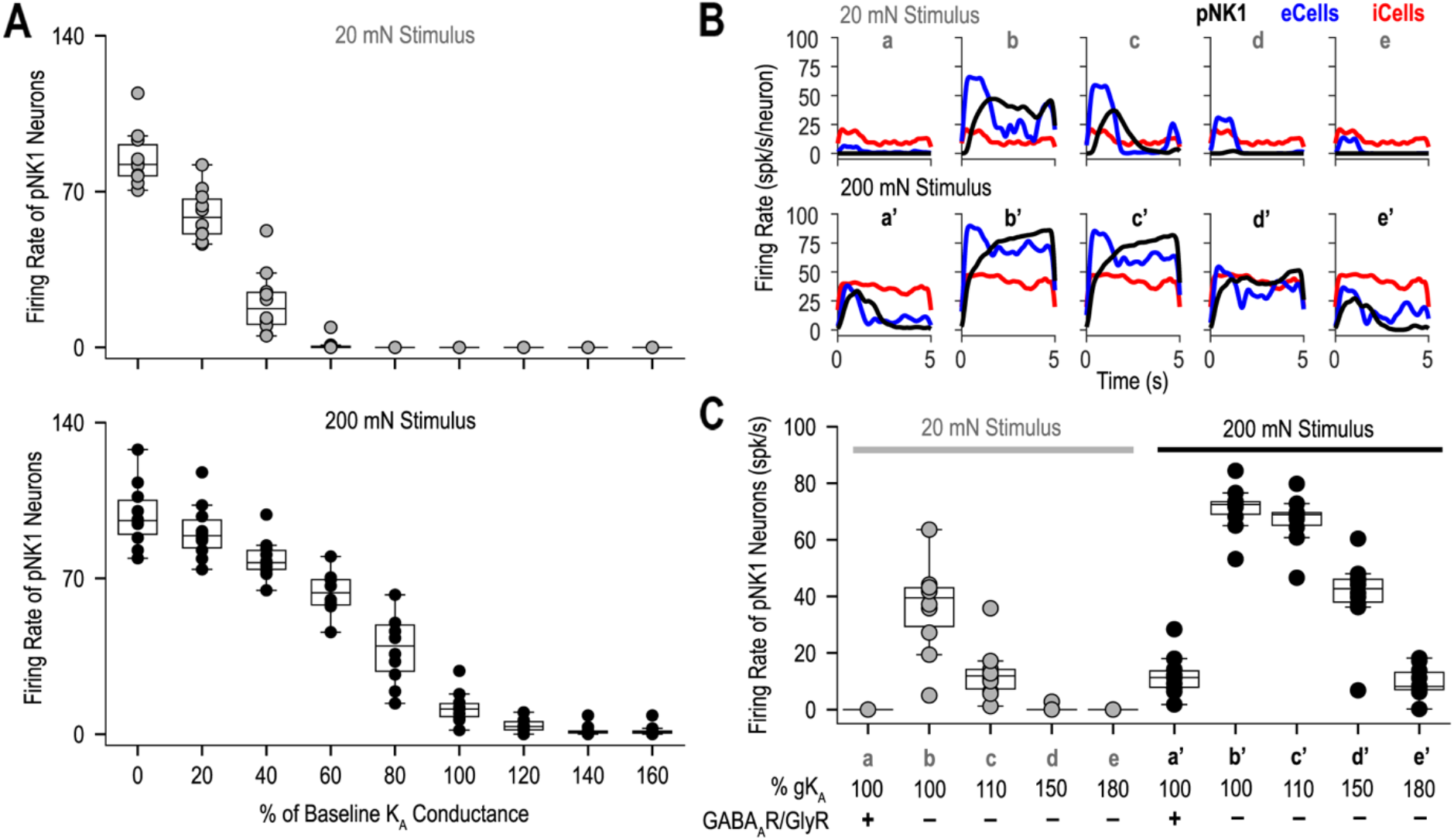
Enhancement of A-type potassium (K_A_) channels can mitigate effects of disinhibition on pNK1 firing. **(A)** Effect of reducing (0-80%) or increasing (120-160%) K_A_ channel density (gKA) on pNK1 firing rates in response to 20 mN (top) or 200 mN (bottom) stimulation. **(B)** Firing rate histograms demonstrating the response of inhibitory (red), excitatory (blue), and projection (black) neurons to 20 and 200 mN (top and bottom, respectively) stimulation across different simulation conditions (see bottom of C). **(C)** Summary of pNK1 firing rates under different simulation conditions. Conditions *a-e* are the same as *a’-e’* but at 20 and 200 mN, respectively. –, blocked; +, intact.

Based on our findings that increased gK_A_ reduces pNK1 firing, we hypothesized that enhancing gK_A_ may rescue the increased firing rates of pNK1 neurons caused by disinhibition. At both 20 and 200 mN complete (100%) blockade of GABA_A_ and glycine receptors resulted in increased firing rates of excitatory interneurons (**Fig. 8B*b,b’***) and pNK1 neurons (**Fig. 8C*b,b’***). While keeping inhibitory receptors blocked, we gradually increased gK_A_ from 100% to 180% of its baseline, which decreased excitatory interneuron firing (blue, **Fig. 8B*c-e,c’-e’***) and, subsequently, pNK1 firing decreased (**Fig. 8C*c-e,c’-e’***). Only a 50% increase in gK_A_ was required to return pNK1 firing rates back to baseline at 20 mN, whereas an 80% increase in gK_A_ was required at 200 mN. These simulation results suggest that decreasing the intrinsic excitability of excitatory interneurons via K_A_ enhancement may be especially effective at mitigating effects of disinhibition during low-threshold stimulation, and thus our model predicts that targeting K_A_ channels might provide a strong therapeutic option against allodynia.

## Discussion

This study presents a new model of the SDH circuit. Synaptic weights were tuned to reproduce pNK1 firing evoked by mechanical stimulation. Different sets of synaptic weights produced similar circuit output under normal conditions, thus revealing solutions to be degenerate. Of the 16 original solutions, one emerged as our top candidate based on its ability to reproduce pNK1 firing rates observed experimentally after disinhibition. Further validation revealed that our top SDH model reproduced expected changes in pNK1 firing after experimental manipulations such as ablation of various neuron types. Apparent inconsistencies – like the relative contribution of iDYN and iPV neurons – were resolved by considering alternative synaptic weight combinations afforded by degeneracy. Our model confirms the role of excitatory interneurons for relaying low-threshold input to projection neurons, and the importance of inhibitory interneurons for gating those circuits.

### Model construction and validation

Accumulated information about spinal neuron spiking properties (**Fig. 1B**) and synaptic connectivity (**Fig. 1D,E**) provided the foundation for our model. We tuned synaptic weights so that pNK1 neuron firing quantitatively reproduced lamina I projection neuron firing rates (Allard, 2019) given primary afferent firing rates (Murthy et al., 2018; Walcher et al., 2018) to comparable mechanical stimuli (**Fig. 2C**). Using a genetic algorithm, we found that many different sets of synaptic weights produce the target output (**Fig. 3A**); in other words, the solution is degenerate (Edelman and Gally, 2001). But only a subset of weight combinations reproduced experimental observations when subsequently tested in other conditions (e.g. disinhibition). To test that our model generalizes to observations not used for tuning, we simulated experimental manipulations shown to modulate projection neuron firing, or are inferred to do so given their impact on pain behavior. Our model quantitatively reproduced the elevated projection neuron firing observed by Allard (2019) after blocking GABA_A_ and glycine receptors (**Fig. 3C*i*** and **6A**). It also reproduced the inferred increase in output caused by ablating iPV neurons and the mitigation of that effect by silencing ePKC*γ* neurons (**Fig. 4**), consistent with Petitjean et al. (2015) and Boyle et al. (2019). Ablating iDYN neurons had little effect on output (**Fig. 5A**), contrary to the large effect reported by Duan et al. (2014), but this can be accounted for by circuit degeneracy (see below).

### Biological insights and recommendations for experiments

Ablation simulations in **Figure 5** predict that inhibition mediated by iPV neurons is more critical for gating low-threshold input to pNK1 neurons than inhibition from iDYN neurons. However, despite targeting different excitatory interneurons, inhibition from iPV and iDYN neurons is seemingly redundant. Weaker iPV-mediated inhibition can be offset by stronger iDYN-mediated inhibition without disrupting SDH output; under those conditions, ablation of iDYN neurons has a larger impact on pNK1 firing (**Fig. 6**), consistent with Duan et al. (2014). Experiments measuring allodynia *and* hyperalgesia (in the same animal) after manipulating iDYN and/or iPV neurons would be useful for reverse engineering the contribution of these populations to the SDH circuit. Generally speaking, measuring multiple different properties/outputs in the same animal provides more information and stronger constraints than measuring each property/output in a different animal (see below). Two circuits that operate equivalently under normal conditions but respond differently to a perturbation, as illustrated by iDYN ablation (**Fig. 6**) and disinhibition (**Fig. 3**), is typical of degenerate systems and can reveal differences in circuit organization (Sakurai et al., 2014; Haddad and Marder, 2018). Such occult differences may explain why two individuals respond differently to the same pathological insult or therapeutic intervention.

In addition to ablating inhibitory interneurons, we also tested the effects of ablating excitatory interneurons (**Fig. 4**). Results confirm the importance of excitatory neurons in forming polysynaptic circuits that relay low-threshold (Aβ) input to pNK1 neurons when not properly gated (Torsney and MacDermott, 2006; Santos et al., 2007; Wang et al., 2013; Duan et al., 2014; Christensen et al., 2016). Our results highlight the multiplicity of subtly different routes through this interconnected circuit. Because of degeneracy, it is difficult to say whether a certain cell type is more important than others – the answer will vary depending on synaptic weights. Here we focus on punctate stimuli, but different types of stimuli (e.g. von Frey vs brush) may preferentially utilize different excitatory routes, making certain routes more or less important for static vs dynamic allodynia (Duan et al., 2014). Furthermore, different chronic pain states (e.g. inflammatory vs neuropathic) likely disrupt the SDH circuit differently (Peirs et al., 2021).

We also investigated the effects of disinhibition (**Fig. 6**) seen under chronic pain conditions. Complete (100%) or partial (90%) inhibitory receptor block and chloride dysregulation allowed pNK1 neurons to be activated by weak stimuli (10-30 mN) as much as they are normally activated by strong stimuli (200 mN), consistent with reduced inhibition causing allodynia (Prescott, 2015). During low-threshold stimulation, chloride dysregulation led to smaller pNK1 firing increases than receptor block, consistent with Keller et al. (2007). Our simulations predict that small differences in disinhibition translate into large differences in pNK1 activation because small effects compound at the circuit level. This could contribute to the differential roles of iPV and iDYN neurons in allodynia and hyperalgesia (see above) and could be explored by optogenetically activating polysynaptic circuits at different points (lamina III vs II) under different inhibitory conditions. Determining whether inhibitory interneurons are activated by excitatory interneurons is also important for deciphering spinal microcircuitry (Prescott and Ratté 2012).

Our simulations offer testable predictions for combating effects of disinhibition. Specifically, our model suggests that blocking NK1 receptors in projection neurons (**Fig. 7**) or reducing intrinsic excitability of excitatory interneurons by modulating K_A_ channels (**Fig. 8**) may mitigate the increased activation of pNK1 neurons following disinhibition. The latter was especially effective at attenuating Aβ input to pNK1 neurons, and is consistent with the established role of spinal K_A_ channels in pain plasticity (Hu et al., 2006, 2007; Zhang et al., 2018) and its potential as a drug target (Wulff et al., 2009; Noh et al., 2019; Yoo et al., 2021). In general, our multiscale model can help predict how a molecular change will manifest at the cellular- or circuit-level. That said, degenerate cells and circuits can adapt, and might do so in ways that subvert therapeutic effects (Ratté et al., 2014; Ratté and Prescott, 2016). Studying such matters requires the computational tools and insights developed in this study.

### Model limitations and outlook for future simulations

Our SDH model remains under-unconstrained because quantitative information about many SDH properties is lacking or indirect. That, together with the high number of free parameters (e.g. synaptic weights), allows for many different parameter sets (solutions) to produce the desired model output. But as reported in **Figure 3**, not all of those solutions produce viable outputs when additional constraints (i.e. disinhibition) are considered. It is important to reserve certain constraints for model validation, rather than applying them during the fitting process. In this respect, the degree of degeneracy could be overestimated and would be revised down in light of additional constraints (Yang et al., 2021). The remaining degeneracy underlies biological variability (see above) and often prevents simple conclusions whose validity hinges on a certain context; in other words, a molecule or cell type may be “necessary” in some circuits but not others contingent on synaptic weights. In general, experiments measuring multiple properties/outputs in the same animal/circuit provide extra information in the form of correlations, which go a long way toward uncovering such contingencies.

We have made certain simplifying assumptions in our model. For instance, intrinsic spiking patterns vary between neuron types and within neurons of a given type (Balachandar and Prescott, 2018), but we only simulated gross differences in pattern between neuron types. Receptive field organization (Hillman and Wall, 1969), descending modulation (Heinricher et al., 2009; Chen and Heinricher, 2019), and presynaptic inhibition (Boyle et al., 2019; Comitato and Bardoni, 2021) are some of the many factors not included in the current model. We focused on punctate mechanical stimulation, but the SDH also processes dynamic mechanical and thermal stimuli, which necessitate consideration of receptive fields to explain spatial and temporal summation (Lee et al., 2019). Such factors may be crucial for resolving differences between static and dynamic allodynia. Furthermore, we have focused on firing rate, but spike timing/patterning (e.g. afferent synchronization) is important for certain types of stimuli like vibration. Such details will be incorporated into future versions of this model and will, we hope, drive new experiments.

In summary, our model reveals how sensory information flows through the SDH circuit, and how that information flow can be pathologically altered in ways that alter pain perception. The effects of pathological disruptions depend on circuit parameters which are, evidently, degenerate. Our model serves as a useful resource that the community is encouraged to use and expand. Integrating additional experimental data into the model will serve not only to strengthen the model, but will also lead to a more cohesive and quantitative understanding of pain processing. This will, we hope, help disentangle how pathological changes in the SDH disrupt pain processing so that robust therapies can be more efficiently developed.

## Notes

### Competing Interest Statement

The authors have declared no competing interest.

## References

Abraira VE et al. (2017) The cellular and synaptic architecture of the mechanosensory dorsal horn. Cell 168:295–310.e19.

Aguiar P, Sousa M, Lima D (2010) NMDA channels together with L-type calcium currents and calcium-activated nonspecific cationic currents are sufficient to generate windup in WDR neurons. J Neurophysiol 104:1155–1166.

Allard J (2019) Physiological properties of the lamina I spinoparabrachial neurons in the mouse. J Physiol 597:2097–2113.

Andrew D, Craig AD (2002) Responses of spinothalamic lamina I neurons to maintained noxious mechanical stimulation in the cat. J Neurophysiol 87:1889–1901.

Arle JE, Carlson KW, Mei L, Iftimia N, Shils JL (2014) Mechanism of dorsal column stimulation to treat neuropathic but not nociceptive pain: analysis with a computational model. Neuromodulation 17:642–655.

Balachandar A, Prescott SA (2018) Origin of heterogeneous spiking patterns from continuously distributed ion channel densities: a computational study in spinal dorsal horn neurons. J Physiol 596:1681–1697.

Bester H, Chapman V, Besson JM, Bernard JF (2000) Physiological properties of the lamina I spinoparabrachial neurons in the rat. J Neurophysiol 83:2239–2259.

Boyle KA, Gradwell MA, Yasaka T, Dickie AC, Polgár E, Ganley RP, Orr DPH, Watanabe M, Abraira VE, Kuehn ED, Zimmermann AL, Ginty DD, Callister RJ, Graham BA, Hughes DI (2019) Defining a spinal microcircuit that gates myelinated afferent input: Implications for tactile allodynia. Cell Rep 28:526–540.e6.

Britton NF, Skevington SM (1989) A mathematical model of the gate control theory of pain. J Theor Biol 137:91–105.

Browne TJ, Smith KM, Gradwell MA, Iredale JA, Dayas CV, Callister RJ, Hughes DI, Graham BA (2019) Spinoparabrachial projection neurons form distinct classes in the mouse dorsal horn. Pain 162:1977–1994.

Cahill CM, Coderre TJ (2002) Attenuation of hyperalgesia in a rat model of neuropathic pain after intrathecal pre- or post-treatment with a neurokinin-1 antagonist. Pain 95:277–285.

Cameron D, Polgár E, Gutierrez-Mecinas M, Gomez-Lima M, Watanabe M, Todd AJ (2015) The organisation of spinoparabrachial neurons in the mouse. Pain 156:2061–2071.

Carlson KD, Nageswaran JM, Dutt N, Krichmar JL (2014) An efficient automated parameter tuning framework for spiking neural networks. Front Neurosci 8:10.

Carnevale NT, Hines ML (2006) The NEURON Book. New York: Cambridge University Press.

Cathenaut L, Leonardon B, Kuster R, Inquimbert P, Schlichter R, Hugel S (2021) Inhibitory interneurons with differential plasticities at their connections tune excitatory/inhibitory balance in the spinal nociceptive system. Pain, in press. doi: 10.1097/j.pain.0000000000002460.

Cheng L, Duan B, Huang T, Zhang Y, Chen Y, Britz O, Garcia-Campmany L, Ren X, Vong L, Lowell BB, Goulding M, Wang Y, Ma Q (2017) Identification of spinal circuits involved in touch-evoked dynamic mechanical pain. Nat Neurosci 20:804–814.

Chen Q, Heinricher MM (2019) Descending control mechanisms and chronic pain. Curr Rheumatol Rep 21:13.

Chien L-Y, Cheng J-K, Chu D, Cheng C-F, Tsaur M-L (2007) Reduced expression of A-type potassium channels in primary sensory neurons induces mechanical hypersensitivity. J Neurosci 27:9855–9865.

Choi S, Hachisuka J, Brett M, Magee A, Koerber H, Ross S, Ginty D (2021) Parallel ascending spinal pathways for affective touch and pain. Nature 587:258–263.

Christensen AJ, Iyer SM, François A, Vyas S, Ramakrishnan C, Vesuna S, Deisseroth K, Scherrer G, Delp SL (2016) In vivo interrogation of spinal mechanosensory circuits. Cell Rep 17:1699–1710.

Comitato A, Bardoni R (2021) Presynaptic inhibition of pain and touch in the spinal cord: from receptors to circuits. Int J Mol Sci 22:414

Coull JAM, Boudreau D, Bachand K, Prescott SA, Nault F, Sík A, De Koninck P, De Koninck Y (2003) Trans-synaptic shift in anion gradient in spinal lamina I neurons as a mechanism of neuropathic pain. Nature 424:938–942.

Crodelle J, Piltz SH, Hagenauer MH, Booth V (2019) Modeling the daily rhythm of human pain processing in the dorsal horn. PLOS Computational Biology 15:e1007106.

Cui L, Kim YR, Kim HY, Lee SC, Shin H-S, Szabó G, Erdélyi F, Kim J, Kim SJ (2011) Modulation of synaptic transmission from primary afferents to spinal substantia gelatinosa neurons by group III mGluRs in GAD65-EGFP transgenic mice. J Neurophysiol 105:1102–1111.

Deuis JR, Dvorakova LS, Vetter I (2017) Methods used to evaluate pain behaviors in rodents. Front Mol Neurosci 10:284.

Duan B, Cheng L, Bourane S, Britz O, Padilla C, Garcia-Campmany L, Krashes M, Knowlton W, Velasquez T, Ren X, Ross S, Lowell BB, Wang Y, Goulding M, Ma Q (2014) Identification of spinal circuits transmitting and gating mechanical pain. Cell 159:1417–1432.

Duan K-Z, Xu Q, Zhang X-M, Zhao Z-Q, Mei Y-A, Zhang Y-Q (2012) Targeting A-type K channels in primary sensory neurons for bone cancer pain in a rat model. Pain 153:562–574.

Dura-Bernal S, Neymotin SA, Kerr CC, Sivagnanam S, Majumdar A, Francis JT, Lytton WW (2017) Evolutionary algorithm optimization of biological learning parameters in a biomimetic neuroprosthesis. IBM J Res Dev 61:6.1–6.14.

Dura-Bernal S, Neymotin SA, Suter BA, Shepherd GMG, Lytton WW (2019a) Multiscale dynamics and information flow in a data-driven model of the primary motor cortex microcircuit. bioRxiv 201707. doi:10.1101/201707.

Dura-Bernal S, Suter BA, Gleeson P, Cantarelli M, Quintana A, Rodriguez F, Kedziora DJ, Chadderdon GL, Kerr CC, Neymotin SA, McDougal RA, Hines M, Shepherd GM, Lytton WW (2019b) NetPyNE, a tool for data-driven multiscale modeling of brain circuits. eLife 8:e44494.

Dura-Bernal S, Zhou X, Neymotin SA, Przekwas A, Francis JT, Lytton WW (2015) Cortical spiking network interfaced with virtual musculoskeletal arm and robotic arm. Front Neurorobot 9:13.

Edelman GM, Gally JA (2001) Degeneracy and complexity in biological systems. Proc Natl Acad Sci U S A 98:13763–13768.

Ferrini F, Perez-Sanchez J, Ferland S, Lorenzo L-E, Godin AG, Plasencia-Fernandez I, Cottet M, Castonguay A, Wang F, Salio C, Doyon N, Merighi A, De Koninck Y (2020) Differential chloride homeostasis in the spinal dorsal horn locally shapes synaptic metaplasticity and modality-specific sensitization. Nat Commun 11:3935.

Gagnon M, Bergeron MJ, Lavertu G, Castonguay A, Tripathy S, Bonin RP, Perez-Sanchez J, Boudreau D, Wang B, Dumas L, Valade I, Bachand K, Jacob-Wagner M, Tardif C, Kianicka I, Isenring P, Attardo G, Coull JAM, De Koninck Y (2013) Chloride extrusion enhancers as novel therapeutics for neurological diseases. Nat Med 19:1524–1528.

Goaillard J-M, Marder E (2021) Ion channel degeneracy, variability, and covariation in neuron and circuit resilience. Annu Rev Neurosci 44:335–357.

Goaillard J-M, Taylor AL, Schulz DJ, Marder E (2009) Functional consequences of animal-to-animal variation in circuit parameters. Nat Neurosci 12:1424–1430.

Gradwell MA, Boyle KA, Browne TJ, Bell AM, Leonardo J, Peralta Reyes FS, Dickie AC, Smith KM, Callister RJ, Dayas CV, Hughes DI, Graham BA (2021) Diversity of inhibitory and excitatory parvalbumin interneuron circuits in the dorsal horn. Pain, in press. doi: 10.1097/j.pain.0000000000002422.

Graham BA, Brichta AM, Callister RJ (2007) Moving from an averaged to specific view of spinal cord pain processing circuits. J Neurophysiol 98:1057–1063.

Grudt TJ, Perl ER (2002) Correlations between neuronal morphology and electrophysiological features in the rodent superficial dorsal horn. J Physiol 540:189–207.

Gutierrez-Mecinas M, Bell AM, Marin A, Taylor R, Boyle KA, Furuta T, Watanabe M, Polgár E, Todd AJ (2017) Preprotachykinin A is expressed by a distinct population of excitatory neurons in the mouse superficial spinal dorsal horn including cells that respond to noxious and pruritic stimuli. Pain 158:440–456.

Gutierrez-Mecinas M, Davis O, Polgár E, Shahzad M, Navarro-Batista K, Furuta T, Watanabe M, Hughes DI, Todd AJ (2019) Expression of calretinin among different neurochemical classes of interneuron in the superficial dorsal horn of the mouse spinal cord. Neuroscience 398:171–181.

Gutierrez-Mecinas M, Furuta T, Watanabe M, Todd AJ (2016) A quantitative study of neurochemically defined excitatory interneuron populations in laminae I-III of the mouse spinal cord. Mol Pain 12:1744806916629065.

Haddad SA, Marder E (2018) Circuit robustness to temperature perturbation is altered by neuromodulators. Neuron 100:609–623.e3.

Hantman AW, van den Pol AN, Perl ER (2004) Morphological and physiological features of a set of spinal substantia gelatinosa neurons defined by green fluorescent protein expression. J Neurosci 24:836–842.

Häring M, Zeisel A, Hochgerner H, Rinwa P, Jakobsson JET, Lönnerberg P, La Manno G, Sharma N, Borgius L, Kiehn O, Lagerström MC, Linnarsson S, Ernfors P (2018) Neuronal atlas of the dorsal horn defines its architecture and links sensory input to transcriptional cell types. Nat Neurosci 21:869–880.

Heinricher MM, Tavares I, Leith JL, Lumb BM (2009) Descending control of nociception: Specificity, recruitment and plasticity. Brain Res Rev 60:214–225.

Hillman P, Wall PD (1969) Inhibitory and excitatory factors influencing the receptive fields of lamina 5 spinal cord cells. Exp Brain Res 9:284–306.

Hines ML, Carnevale NT (2001) NEURON: a tool for neuroscientists. Neuroscientist 7:123–135.

Huang H-Y, Cheng J-K, Shih Y-H, Chen P-H, Wang C-L, Tsaur M-L (2005) Expression of A-type K channel alpha subunits Kv4.2 and Kv4.3 in rat spinal lamina II excitatory interneurons and colocalization with pain-modulating molecules. Eur J Neurosci 22:1149–1157.

Huang J, Polgár E, Solinski HJ, Mishra SK, Tseng P-Y, Iwagaki N, Boyle KA, Dickie AC, Kriegbaum MC, Wildner H, Zeilhofer HU, Watanabe M, Riddell JS, Todd AJ, Hoon MA (2018) Circuit dissection of the role of somatostatin in itch and pain. Nat Neurosci 21:707–716.

Hu H-J, Alter BJ, Carrasquillo Y, Qiu C-S, Gereau RW (2007) Metabotropic glutamate receptor 5 modulates nociceptive plasticity via extracellular signal-regulated kinase–Kv4.2 signaling in spinal cord dorsal horn neurons. J Neurosci 27:13181–13191.

Hu H-J, Carrasquillo Y, Karim F, Jung WE, Nerbonne JM, Schwarz TL, Gereau RW 4th (2006) The Kv4.2 potassium channel subunit is required for pain plasticity. Neuron 50:89–100.

Hunt CA, Erdemir A, Lytton WW, Mac Gabhann F, Sander EA, Transtrum MK, Mulugeta L (2018) The spectrum of mechanism-oriented models and methods for explanations of biological phenomena. Processes 6:56.

Jensen TS, Finnerup NB (2014) Allodynia and hyperalgesia in neuropathic pain: clinical manifestations and mechanisms. Lancet Neurol 13:924–935.

Kardon AP et al. (2014) Dynorphin acts as a neuromodulator to inhibit itch in the dorsal horn of the spinal cord. Neuron 82:573–586.

Keller AF, Beggs S, Salter MW, De Koninck Y (2007) Transformation of the output of spinal lamina I neurons after nerve injury and microglia stimulation underlying neuropathic pain. Mol Pain 3:27.

Le Bars D, Gozariu M, Cadden SW (2001) Animal models of nociception. Pharmacol Rev 53:597–652.

Lechner SG (2017) An update on the spinal and peripheral pathways of pain signalling. e-Neuroforum 23:A131–A136.

Lee KY, Prescott SA (2015) Chloride dysregulation and inhibitory receptor blockade yield equivalent disinhibition of spinal neurons yet are differentially reversed by carbonic anhydrase blockade. Pain 156:2431–2437.

Lee KY, Ratté S, Prescott SA (2019) Excitatory neurons are more disinhibited than inhibitory neurons by chloride dysregulation in the spinal dorsal horn. eLife 8:e49753.

Le Franc Y, Le Masson G (2010) Multiple firing patterns in deep dorsal horn neurons of the spinal cord: Computational analysis of mechanisms and functional implications. J Neurophysiol 104:1978–1996.

Lu Y, Dong H, Gao Y, Gong Y, Ren Y, Gu N, Zhou S, Xia N, Sun Y-Y, Ji R-R, Xiong L (2013) A feed-forward spinal cord glycinergic neural circuit gates mechanical allodynia. J Clin Invest 123:4050–4062.

Lu Y, Perl ER (2003) A specific inhibitory pathway between substantia gelatinosa neurons receiving direct C-fiber input. J Neurosci 23:8752–8758.

Lu Y, Perl ER (2005) Modular organization of excitatory circuits between neurons of the spinal superficial dorsal horn (laminae I and II). J Neurosci 25:3900–3907.

Lytton WW, Arle J, Bobashev G, Ji S, Klassen TL, Marmarelis VZ, Schwaber J, Sherif MA, Sanger TD (2017) Multiscale modeling in the clinic: diseases of the brain and nervous system. Brain Inform 4:219–230.

Mantyh PW, Rogers SD, Honore P, Allen BJ, Ghilardi JR, Li J, Daughters RS, Lappi DA, Wiley RG, Simone DA (1997) Inhibition of hyperalgesia by ablation of lamina I spinal neurons expressing the substance P receptor. Science 278:275–279.

Mapplebeck JCS, Lorenzo L-E, Lee KY, Gauthier C, Muley MM, De Koninck Y, Prescott SA, Salter MW (2019) Chloride dysregulation through downregulation of KCC2 mediates neuropathic pain in both sexes. Cell Rep 28:590–596.e4.

Marder E (2011) Variability, compensation, and modulation in neurons and circuits. Proc Natl Acad Sci USA 108 Suppl 3:15542–15548.

Marder E, Goeritz ML, Otopalik AG (2015) Robust circuit rhythms in small circuits arise from variable circuit components and mechanisms. Curr Opin Neurobiol 31:156–163.

Melnick IV, Santos SFA, Szokol K, Szûcs P, Safronov BV (2004) Ionic basis of tonic firing in spinal substantia gelatinosa neurons of rat. J Neurophysiol 91:646–655.

Melzack R, Wall PD (1965) Pain mechanisms: a new theory. Science 150:971–979.

Murthy SE, Loud MC, Daou I, Marshall KL, Schwaller F, Kühnemund J, Francisco AG, Keenan WT, Dubin AE, Lewin GR, Patapoutian A (2018) The mechanosensitive ion channel Piezo2 mediates sensitivity to mechanical pain in mice. Sci Transl Med 10:eaat9897.

Neymotin SA, Suter BA, Dura-Bernal S, Shepherd GMG, Migliore M, Lytton WW (2017) Optimizing computer models of corticospinal neurons to replicate in vitro dynamics. J Neurophysiol 117:148–162.

Nichols ML, Allen BJ, Rogers SD, Ghilardi JR, Honore P, Luger NM, Finke MP, Li J, Lappi DA, Simone DA, Mantyh PW (1999) Transmission of chronic nociception by spinal neurons expressing the substance P receptor. Science 286:1558–1561.

Noh W, Pak S, Choi G, Yang S, Yang S (2019) Transient potassium channels: therapeutic targets for brain disorders. Front Cell Neurosci 13:265.

O’Leary T, Sutton AC, Marder E (2015) Computational models in the age of large datasets. Curr Opin Neurobiol 32:87–94.

Peirs C, Dallel R, Todd AJ (2020) Recent advances in our understanding of the organization of dorsal horn neuron populations and their contribution to cutaneous mechanical allodynia. J Neural Transm 127:505–525.

Peirs C, Seal RP (2016) Neural circuits for pain: recent advances and current views. Science 354:578–584.

Peirs C, Williams S-PG, Zhao X, Arokiaraj CM, Ferreira DW, Noh M-C, Smith KM, Halder P, Corrigan KA, Gedeon JY, Lee SJ, Gatto G, Chi D, Ross SE, Goulding M, Seal RP (2021) Mechanical allodynia circuitry in the dorsal horn is defined by the nature of the injury. Neuron 109:73–90.e7.

Peirs C, Williams S-PG, Zhao X, Walsh CE, Gedeon JY, Cagle NE, Goldring AC, Hioki H, Liu Z, Marell PS, Seal RP (2015) Dorsal horn circuits for persistent mechanical pain. Neuron 87:797–812.

Petitjean H, Bourojeni FB, Tsao D, Davidova A, Sotocinal SG, Mogil JS, Kania A, Sharif-Naeini R (2019) Recruitment of spinoparabrachial neurons by dorsal horn calretinin neurons. Cell Rep 28:1429–1438.e4.

Petitjean H, Pawlowski SA, Fraine SL, Sharif B, Hamad D, Fatima T, Berg J, Brown CM, Jan L-Y, Ribeiro-da-Silva A, Braz JM, Basbaum AI, Sharif-Naeini R (2015) Dorsal horn parvalbumin neurons are gate-keepers of touch-evoked pain after nerve injury. Cell Rep 13:1246–1257.

Peyronnard JM, Charron LF, Lavoie J, Messier JP (1986) Motor, sympathetic and sensory innervation of rat skeletal muscles. Brain Res 373:288–302.

Polgár E, Durrieux C, Hughes DI, Todd AJ (2013) A quantitative study of inhibitory interneurons in laminae I-III of the mouse spinal dorsal horn. PLoS One 8:e78309.

Polgár E, Hughes DI, Riddell JS, Maxwell DJ, Puskár Z, Todd AJ (2003) Selective loss of spinal GABAergic or glycinergic neurons is not necessary for development of thermal hyperalgesia in the chronic constriction injury model of neuropathic pain. Pain 104:229–239.

Prescott SA (2015) Synaptic inhibition and disinhibition in the spinal dorsal horn. In: Progress in Molecular Biology and Translational Science (Price TJ, Dussor G, eds), pp 359–383. Academic Press.

Prescott SA, De Koninck Y (2002) Four cell types with distinctive membrane properties and morphologies in lamina I of the spinal dorsal horn of the adult rat. J Physiol 539:817–836.

Prescott SA, Ma Q, De Koninck Y (2014) Normal and abnormal coding of somatosensory stimuli causing pain. Nat Neurosci 17:183–191.

Prescott SA, Sejnowski TJ, De Koninck Y (2006) Reduction of anion reversal potential subverts the inhibitory control of firing rate in spinal lamina I neurons: towards a biophysical basis for neuropathic pain. Mol Pain 2:32.

Prinz AA, Bucher D, Marder E (2004) Similar network activity from disparate circuit parameters. Nat Neurosci 7:1345–1352.

Punnakkal P, von Schoultz C, Haenraets K, Wildner H, Zeilhofer HU (2014) Morphological, biophysical and synaptic properties of glutamatergic neurons of the mouse spinal dorsal horn. J Physiol 592:759–776.

Ratté S, Lankarany M, Rho Y-A, Patterson A, Prescott SA (2015) Subthreshold membrane currents confer distinct tuning properties that enable neurons to encode the integral or derivative of their input. Front Cell Neurosci 8:452.

Ratté S, Prescott SA (2015) Pain processing by spinal microcircuits: afferent combinatorics. Curr Opin Neurobiol 22:631–639.

Ratté S, Prescott SA (2016) Afferent hyperexcitability in neuropathic pain and the inconvenient truth about its degeneracy. Curr Opin Neurobiol 36:31–37.

Ratté S, Zhu Y, Lee KY, Prescott SA (2014) Criticality and degeneracy in injury-induced changes in primary afferent excitability and the implications for neuropathic pain. eLife 3:e02370.

Ruscheweyh R, Sandkühler J (2002) Lamina-specific membrane and discharge properties of rat spinal dorsal horn neurones in vitro. J Physiol 541:231–244.

Sakurai A, Tamvacakis AN, Katz PS (2014) Hidden synaptic differences in a neural circuit underlie differential behavioral susceptibility to a neural injury. eLife 3:e02598.

Santos SFA, Rebelo S, Derkach VA, Safronov BV (2007) Excitatory interneurons dominate sensory processing in the spinal substantia gelatinosa of rat. J Physiol 581:241–254.

Shimazaki H, Shinomoto S (2010) Kernel bandwidth optimization in spike rate estimation. J Comput Neurosci 29:171–182.

Sivagnanam S, Gorman W, Doherty D, Neymotin SA, Fang S, Hovhannisyan H, Lytton WW, Dura-Bernal S (2020) Simulating large-scale models of brain neuronal circuits using Google Cloud Platform. Practice and Experience in Advanced Research Computing. Association for Computing Machinery, New York, NY, USA, 505–509.

Smith KM, Browne TJ, Davis OC, Coyle A, Boyle KA, Watanabe M, Dickinson SA, Iredale JA, Gradwell MA, Jobling P, Callister RJ, Dayas CV, Hughes DI, Graham BA (2019) Calretinin positive neurons form an excitatory amplifier network in the spinal cord dorsal horn. eLife 8:e49190.

Tang LS, Taylor AL, Rinberg A, Marder E (2012) Robustness of a rhythmic circuit to short- and long-term temperature changes. Journal of Neuroscience 32:10075–10085.

Todd AJ (2010) Neuronal circuitry for pain processing in the dorsal horn. Nat Rev Neurosci 11:823–836.

Todd AJ (2017) Identifying functional populations among the interneurons in laminae I-III of the spinal dorsal horn. Mol Pain 13:1744806917693003.

Todd AJ, McGill MM, Shehab SA (2000) Neurokinin 1 receptor expression by neurons in laminae I, III and IV of the rat spinal dorsal horn that project to the brainstem. Eur J Neurosci 12:689–700.

Torsney C, MacDermott AB (2006) Disinhibition opens the gate to pathological pain signaling in superficial neurokinin 1 receptor-expressing neurons in rat spinal cord. J Neurosci 26:1833–1843.

Walcher J, Ojeda-Alonso J, Haseleu J, Oosthuizen MK, Rowe AH, Bennett NC, Lewin GR (2018) Specialized mechanoreceptor systems in rodent glabrous skin. J Physiol 596:4995–5016.

Wang D, Tawfik VL, Corder G, Low SA, François A, Basbaum AI, Scherrer G (2018) Functional divergence of delta and mu opioid receptor organization in CNS pain circuits. Neuron 98:90–108.e5.

Wang X, Zhang J, Eberhart D, Urban R, Meda K, Solorzano C, Yamanaka H, Rice D, Basbaum AI (2013) Excitatory superficial dorsal horn interneurons are functionally heterogeneous and required for the full behavioral expression of pain and itch. Neuron 78:312–324.

Wulff H, Castle NA, Pardo LA (2009) Voltage-gated potassium channels as therapeutic targets. Nat Rev Drug Discov 8:982–1001.

Yang J, Shakil H, Ratté S, Prescott SA (2021) Co-regulating many properties by co-adjusting few ion channels increases ion channel correlations and the risk of regulation failure. bioRxiv 410787. doi:2020.12.04.410787.

Yasaka T, Tiong SYX, Hughes DI, Riddell JS, Todd AJ (2010) Populations of inhibitory and excitatory interneurons in lamina II of the adult rat spinal dorsal horn revealed by a combined electrophysiological and anatomical approach. Pain 151:475–488.

Yekkirala AS, Roberson DP, Bean BP, Woolf CJ (2017) Breaking barriers to novel analgesic drug development. Nat Rev Drug Discov 16:545–564.

Yoo S, Santos C, Reynders A, Marics I, Malapert P, Gaillard S, Charron A, Ugolini S, Rossignol R, El Khallouqi A, Springael J-Y, Parmentier M, Saurin AJ, Goaillard J-M, Castets F, Clerc N, Moqrich A (2021) TAFA4 relieves injury-induced mechanical hypersensitivity through LDL receptors and modulation of spinal A-type K+ current. Cell Rep 37:109884.

Zeilhofer HU, Wildner H, Yévenes GE (2012) Fast synaptic inhibition in spinal sensory processing and pain control. Physiol Rev 92:193–235.

Zhang TC, Janik JJ, Grill WM (2014) Modeling effects of spinal cord stimulation on wide-dynamic range dorsal horn neurons: influence of stimulation frequency and GABAergic inhibition. J Neurophysiol 112:552–567.

Zhang TC, Janik JJ, Peters RV, Chen G, Ji R-R, Grill WM (2015) Spinal sensory projection neuron responses to spinal cord stimulation are mediated by circuits beyond gate control. J Neurophysiol 114:284–300.

Zhang Y, Liu S, Zhang Y-Q, Goulding M, Wang Y-Q, Ma Q (2018) Timing mechanisms underlying gate control by feedforward inhibition. Neuron 99:941–955.e4.

